# Enhanced excitability of the hippocampal CA2 region and its contribution to seizure generation in a mouse model of temporal lobe epilepsy

**DOI:** 10.1101/2022.02.02.478736

**Authors:** Alexander C. Whitebirch, John J. LaFrancois, Swati Jain, Paige Leary, Bina Santoro, Steven A. Siegelbaum, Helen E. Scharfman

## Abstract

The hippocampal CA2 region, an area important for social memory, has been suspected to play a role in temporal lobe epilepsy (TLE) because of its resistance to the degeneration observed in neighboring CA1 and CA3 regions in both human and rodent models of TLE. However, little is known about whether alterations in CA2 properties serve to promote seizure generation or propagation. Here we have used the pilocarpine-induced status epilepticus (PILO-SE) model of TLE to explore the role of CA2. *Ex vivo* electrophysiological recordings from acute hippocampal slices revealed a set of coordinated changes that enhance CA2 intrinsic excitability, reduce CA2 local inhibitory input, and increase CA2 excitatory output to its major CA1 synaptic target. Moreover, selective silencing of CA2 pyramidal cells using a chemogenetic approach caused a significant decrease in the frequency of spontaneous seizures. These findings provide the first evidence that CA2 actively contributes to TLE seizure activity and may thus be a promising therapeutic target.

## INTRODUCTION

Temporal lobe epilepsy (TLE) is among the most prevalent neurological disorders, with approximately one third of patients experiencing seizures that are refractory to medication (P. Kwan & Sander, 2004; Patrick Kwan & Brodie, 2000). The critical task of identifying new therapeutic targets in TLE therefore requires new insight into the mechanisms of seizure generation. Anatomical and functional studies have suggested that the relatively unexplored CA2 region of the hippocampus may play an important role in seizure generation. Thus, autopsy specimens from human TLE patients display a characteristic pattern of hippocampal neurodegeneration termed mesial temporal sclerosis (MTS), with a substantial loss of neurons in the hilus of the dentate gyrus hilus and in the CA3 and CA1 pyramidal cell layer. In contrast, there is a relative sparing of dentate gyrus granule cells (GCs) and CA2 pyramidal cells (PCs) (Blümcke et al., 2013; Steve, Jirsch, & Gross, 2014).

Although relatively small, the CA2 subfield is of particular interest as it forms the nexus of a powerfully excitatory disynaptic circuit that directly links cortical input to hippocampal output (Chevaleyre & Siegelbaum, 2010; Srinivas et al., 2017). Furthermore, accumulating evidence suggests that CA2 may have a unique role regulating hippocampal network excitability and organizing population activity (Boehringer et al., 2017; He et al., 2021; Lehr et al., 2021; Oliva, Fernández-Ruiz, Buzsáki, & Berényi, 2016; Oliva, Fernández-Ruiz, Leroy, & Siegelbaum, 2020). Recordings from surgically-resected hippocampal tissue from patients with refractory TLE revealed spontaneous interictal-like spikes in CA2, but the study could not address whether CA2 activity contributes to spontaneous recurring seizures (Wittner et al., 2009). Electrophysiological studies utilizing human tissue are also limited in that healthy control tissue is not available. Examination of both surgically-resected human epileptic tissue and rodent models of TLE has revealed alterations to the CA2 subfield, including synaptic reorganization of dentate gyrus granule cell mossy fiber axons in CA2 (Freiman et al., 2021; Häussler, Rinas, Kilias, Egert, & Haas, 2016) and decreases in parvalbumin expression (Andrioli & Arellano, 2007; Wittner et al., 2009). Altogether, this evidence suggests that CA2 may be an important component of a seizure-generating epileptic network in TLE. To date, however, there has been no direct test of the hypothesis that CA2 contributes to seizure generation in chronic epilepsy.

Here we tested that hypothesis using the pilocarpine-induced status epilepticus (PILO-SE) model of TLE. We took advantage of the Amigo2-Cre mouse line which enabled the relatively selective targeting of CA2 using Cre-dependent viral vectors (Hitti & Siegelbaum, 2014). Using this approach we expressed chemogenetic and optogenetic probes in CA2 pyramidal cells (PCs) and found that in SE mice with spontaneous recurrent seizures chemogenetic silencing of CA2 significantly reduced seizure frequency. Moreover, *ex vivo* recordings from acute hippocampal slices revealed that PILO-SE enhanced the intrinsic excitability of surviving CA2 neurons, decreased CA2 synaptic inhibition, increased synaptic excitation of CA2 by its mossy fiber inputs, and increased CA2 excitatory output onto its downstream CA1 targets. Thus, our results point to an important role of CA2 in seizure generation and/or propagation that makes it an attractive target for novel therapeutic interventions.

## RESULTS

Are there alterations in CA2 intrinsic excitability or extrinsic synaptic properties associated with PILO-SE that may contribute to the emergence or propagation of seizures in TLE? To address this question, we performed whole-cell recordings from PCs in acute hippocampal slices prepared from SE mice and age-matched controls. We first examined whether SE produced changes in CA2 PC intrinsic excitability. Next we addressed whether the TLE model altered excitatory or inhibitory synaptic responses of CA2 PCs upon activation of their major synaptic input pathways. We then examined whether TLE modified the strength of CA2 PC synaptic output to its major target populations. Finally, we examined *in vivo* whether CA2 contributed to seizure frequency, duration, or severity in SE animals.

### PILO-SE treatment enhances CA2 intrinsic excitability

When we injected depolarizing current steps to elicit action potentials (one-second square pulses ranging from 100 pA to 1000 pA), we found a significant increase in excitability of CA2 PCs from SE mice compared to control animals. Thus, the firing rate versus current curve was shifted to higher firing rates for a given depolarizing current injection in CA2 PCs from SE mice relative to controls (figure 1B, C; two-way ANOVA with Sidak’s test; \*\*\*\**P* < 0.0001; n = 95 control, 138 SE). The maximum firing rate achieved in each cell over the course of the ten current steps was also significantly higher in cells from SE mice compared to controls (figure 1D; Mann-Whitney; SE vs control, \*\*\*\**P* < 0.0001; n = 92 control, 138 SE).

**Figure 1.**
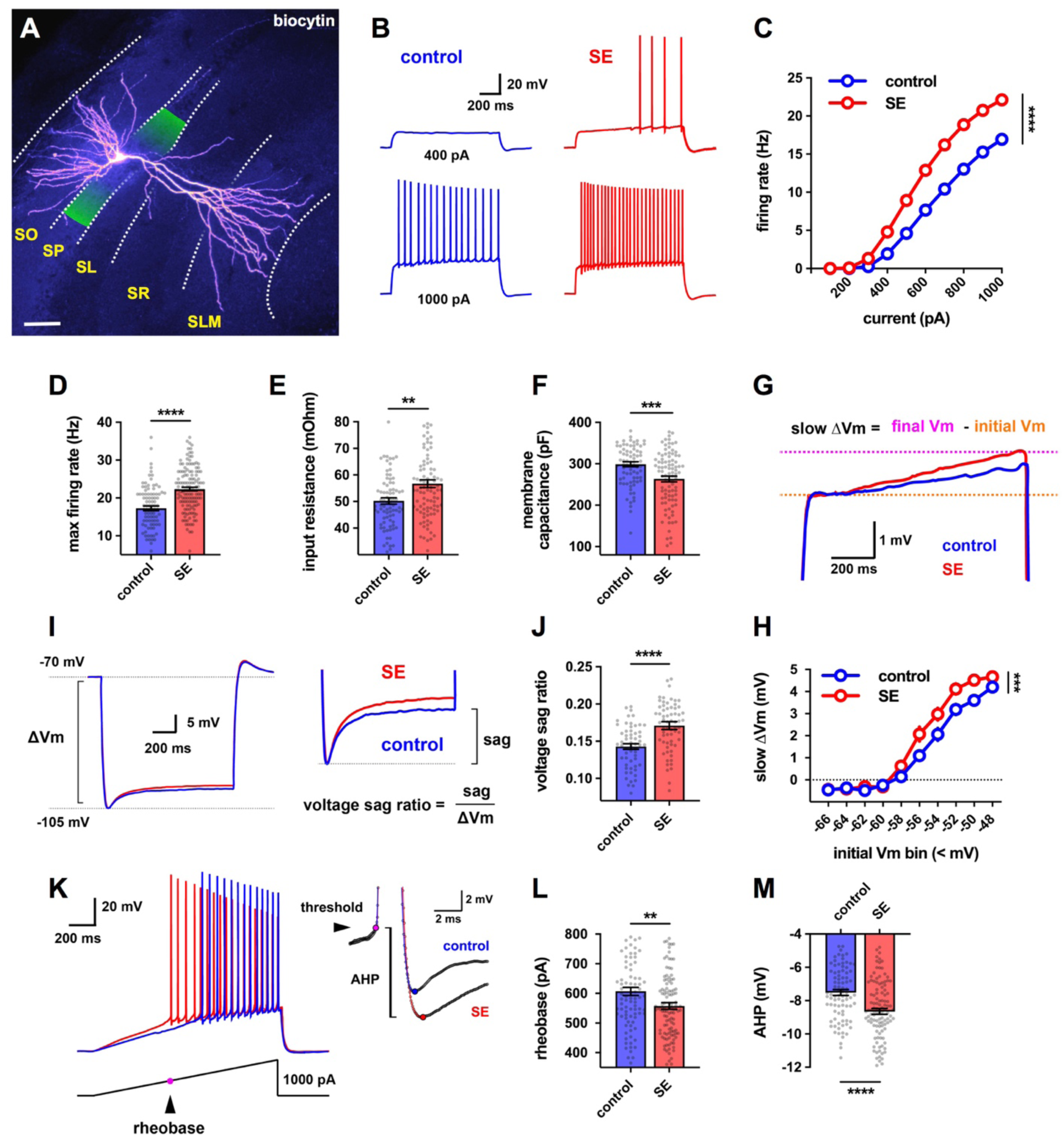
CA2 pyramidal cells (PCs) in slices from pilocarpine-treated mice had increased intrinsic excitability. **(A)** A CA2 PC labeled with intracellular biocytin, with the approximate bounds of the CA2 subfield shaded in green. The hippocampal layers in this and all subsequent images are labeled as follows: stratum oriens (SO), stratum pyramidale (SP), stratum lucidum (SL), stratum radiatum (SR), stratum lacunosum moleculare (SLM). Scale bar is 60 μm. **(B)** Representative traces of membrane depolarization and action potential firing patterns in control (blue) and SE (red) CA2 PCs in response to a 1 second current step. **(C)** CA2 PCs in slices from SE mice fire a greater number of action potentials in response to depolarizing current steps. **(D)** The maximum firing rate was increased in CA2 PCs from SE mice. **(E)** Input resistance was increased in CA2 PCs from SE mice. **(F)** Membrane capacitance was reduced in CA2 PCs from SE mice. **(G)** Representative averaged traces of the slow ramping depolarization exhibited by CA2 PCs near action potential threshold. **(H)** Cells in slices from SE mice exhibited a larger slow depolarization. **(I)** Representative averaged traces from control and SE CA2 PCs showing hyperpolarization-induced membrane voltage sag. **(J)** The voltage sag ratio was greater in CA2 PCs from SE mice. **(K)** Representative membrane voltage responses from control (blue) and SE (red) CA2 PCs to a ramp of applied current. **(L)** The rheobase current was reduced in CA2 PCs from SE mice. **(M)** The amplitude of the AHP was significantly increased in CA2 PCs from SE mice.

We found no difference in resting membrane potential between CA2 PCs from control or SE mice (Mann-Whitney; *P* = 0.3915; n = 119 control, 143 SE). In contrast, we saw a significant increase in input resistance (R_in_) in cells from SE mice relative to controls (figure 1E; Mann-Whitney; \*\**P* = 0.0022; n = 81 control, 94 SE). We also observed a reduction in membrane capacitance (Cm) in cells from SE mice (figure 1F; Mann-Whitney; \*\*\**P* = 0.0005; n = 78 control, 92 SE), suggesting a decreased membrane surface area. In addition, a slow ramping depolarization evident during current steps that is characteristic of CA2 PCs (Kohara et al., 2014) was increased in amplitude in CA2 PCs from SE mice relative to controls (figure 1G, H; mixed-effects model with Holm-Sidak’s test; ***P = 0.0001; n = 110 control, 129 cells from SE mice). Furthermore, we saw an increase in voltage sag in response to hyperpolarizing current steps (figure 1I, J; Mann-Whitney; ****P < 0.0001; n = 57 control, 67 SE), indicative of an increased hyperpolarization-activated HCN channel current (Srinivas et al., 2017).

We next examined action potential features in more detail by applying a 1-second depolarizing current ramp (figure 1K). The minimal current needed to elicit an action potential (rheobase) was significantly reduced in cells from SE mice (figure 1L; Mann-Whitney; **P = 0.0138; n = 88 control, 121 SE), consistent with the increased action potential firing during the current steps. However, there was no change in action potential voltage threshold, amplitude, half-width, maximum rate of rise, and maximum rate of descent, suggesting that the increase in spike firing may be due to the increase in input resistance, rather than to a change in voltage-gated channels. Although action potential parameters were largely unchanged, the fast afterhyperpolarization (AHP) was significantly larger in amplitude in CA2 PCs from SE animals relative to cells from control mice (figure 1K, M; Mann-Whitney; ****P < 0.0001; n = 88 control, 120 SE). Measures of intrinsic electrophysiological properties are summarized in supplementary table 1.

### PILO-SE reduces synaptic inhibition but not excitation of CA2 PCs by their CA3 inputs, with no change in synaptic response to CA2 cortical inputs

CA2 receives its major excitatory inputs from the entorhinal cortical perforant path axons and CA3 PC Schaffer collaterals (Chevaleyre & Siegelbaum, 2010). In addition, CA2 receives recurrent collateral excitatory synapses from other CA2 neurons and weaker excitatory input from the mossy fibers of dentate gyrus granule cells (Kohara et al., 2014; Okamoto & Ikegaya, 2018). These excitatory inputs also recruit strong inhibition of CA2 PCs, mediated by a diverse and widespread population of GABAergic interneurons (Nasrallah et al., 2019; Sun et al., 2017). We next systematically evaluated whether the PILO-SE model of TLE affects the strength of these excitatory and inhibitory synaptic connections. We first examined the effects of electrical stimulation of local synaptic inputs using a stimulating electrode in the nearby stratum radiatum (SR), which is dominated by excitatory Schaffer collateral (SC) inputs and inhibitory inputs from local interneurons (figure 2A).

**Figure 2.**
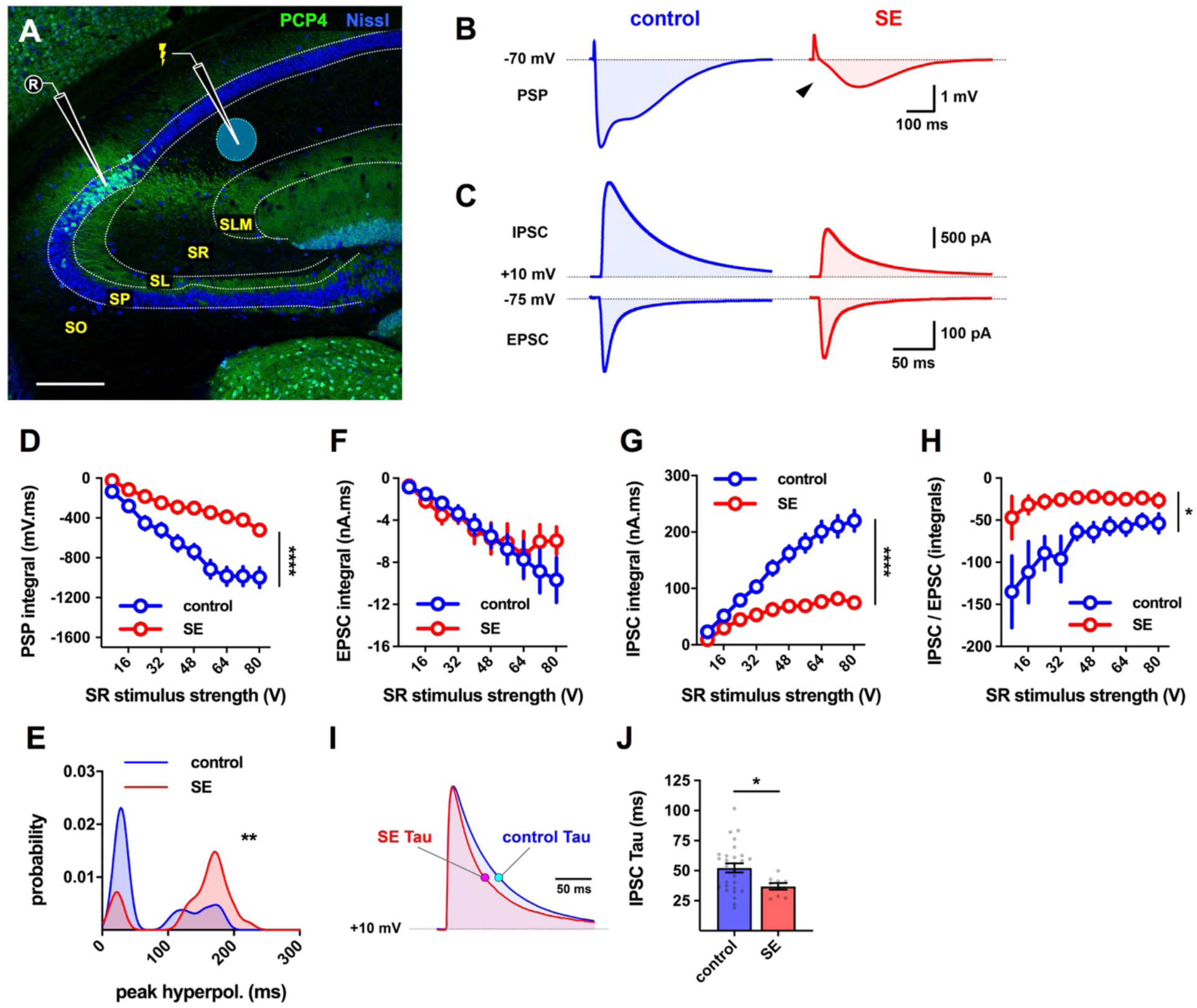
Inhibition of CA2 PNs recruited by stimulation of CA2/CA3 axons was diminished in slices from SE mice. **(A)** Representative hippocampal section stained for Nissl (blue) and PCP4 (green) to illustrate the configuration used to measure synaptic input to CA2 PCs from the CA2/3 local collaterals in the SR. Scale bar is 250 μm. **(B)** Representative averaged PSPs from control and SE CA2 PCs in response to SR stimulation. **(C)** Top, representative averaged SR-evoked inhibitory postsynaptic currents (IPSCs) recorded in voltage clamp configuration in the presence of intracellular cesium (*Cs+*) from CA2 PCs voltage-clamped at +10 mV. Below, SR-evoked excitatory postsynaptic currents (EPSC) from CA2 PCs voltage-clamped at -75 mV. **(D)** The integral of the SR-evoked postsynaptic potential was significantly less negative in CA2 PCs from SE mice. **(E)** Probability density functions constructed from the measured latencies between stimulation and the peak hyperpolarization of SR-evoked PSPs. **(F - H)** Input-output curves of the integral of the SR-evoked EPSC (F), the integral of the SR-evoked IPSC (G), and the ratio between the integrals of the IPSC and EPSC (H). **(I)** Representative averaged traces illustrating the time course of SR-evoked IPSCs with the exponential time constant of decay, tau, indicated for the currents from cells from control and SE mice. **(J)** The time constant of the SR-evoked IPSC was faster in CA2 PCs from SE mice.

In current clamp recordings from CA2 PCs from control mice, SR stimulation evoked a triphasic postsynaptic potential (PSP) consisting of a small initial depolarization followed by a larger hyperpolarization (figure 2B, left). In CA2 PCs from epileptic mice we observed a striking decrease in the magnitude of the prolonged hyperpolarization (figure 2B, right). As a result, the net integral of the membrane voltage (Vm) response (figure 2D) was significantly smaller in cells from SE mice compared to controls, suggesting a reduction in inhibition (mixed-effects model with Holm-Sidak’s test; \*\*\*\**P* < 0.0001; n = 36 control, 35 SE).

The triphasic PSP observed in CA2 PCs reflects the summation of both monosynaptic excitation, from CA2 and CA3 associational collaterals, and inhibition from interneurons that consists of both fast and slow hyperpolarizing components mediated by GABA_A_ and GABA_B_ receptors respectively. We observed a pronounced loss of fast inhibition in SE mice relative to controls (figure 2B, arrowhead). This altered the dynamics of synaptic inhibition, resulting in a significant shift in the distribution of latency to peak inhibition to longer times (figure 2E; Kolmogorov-Smirnov; \*\**P* = 0.0018; n = 25 control, 30 SE).

We next performed voltage clamp recordings of isolated SR-evoked excitatory and inhibitory fast postsynaptic currents (EPSCs and IPSCs, respectively). When we recorded the IPSC by voltage-clamping the membrane near the reversal potential of the EPSC (+10 mV), we observed a marked reduction in the IPSC amplitude compared to that in cells from control mice. However, we saw little change in the EPSC amplitude relative to control values when the membrane was voltage clamped near the IPSC reversal potential (−75 mV) (figure 2C). Measurement of the integral of the synaptic currents (figure 2F, G) confirmed a significant decrease in the total charge carried by the IPSC (mixed-effects model; *P* < 0.0001; n = 32 control, 15 SE) with no change in the EPSC charge (mixed-effects model; *P* = 0.7860; n = 31 control, 15 SE), resulting in a pronounced decrease in the IPSC/EPSC ratio (figure 2H; mixed-effects model; \**P* = 0.0389; n = 30 controls, 14 SE). We also found that the time course of IPSC decay was significantly faster in CA2 PCs from SE mice compared to controls, as quantified by a decrease in the exponential decay time constant (figure 2I, J, Mann-Whitney; \**P* = 0.0403; n = 28 controls, 9 cells from SE mice). In contrast to the reduced synaptic inhibition in response to electrical stimulation of SR inputs, PILO-SE caused no significant difference, relative to control values, in either synaptic inhibition or excitation that was evoked by the direct entorhinal cortical projections to CA2 using a stimulating electrode in stratum lacunosum moleculare (SLM) (supp. figure 2).

### PILO-SE reduces inhibition but not excitation evoked by optogenetic activation of CA2 recurrent collaterals

Next we examined whether PILO-SE altered synaptic responses evoked by optogenetic activation of the CA2 PC recurrent connections. We stereotactically injected adenoassociated virus (AAV) in the dorsal hippocampus of Amigo2-Cre mice to drive expression of channelrhodopsin-2 (ChR2-eYFP) selectively in CA2 PCs (see methods, supp. figure 2). In acute hippocampal slices photostimulation (2 ms pulses) effectively triggered CA2 action potential output (supp. figure 2B), which evoked mixed excitatory-inhibitory synaptic responses recorded from CA2 PCs not expressing ChR2 (figure 3A - C).

**Figure 3.**
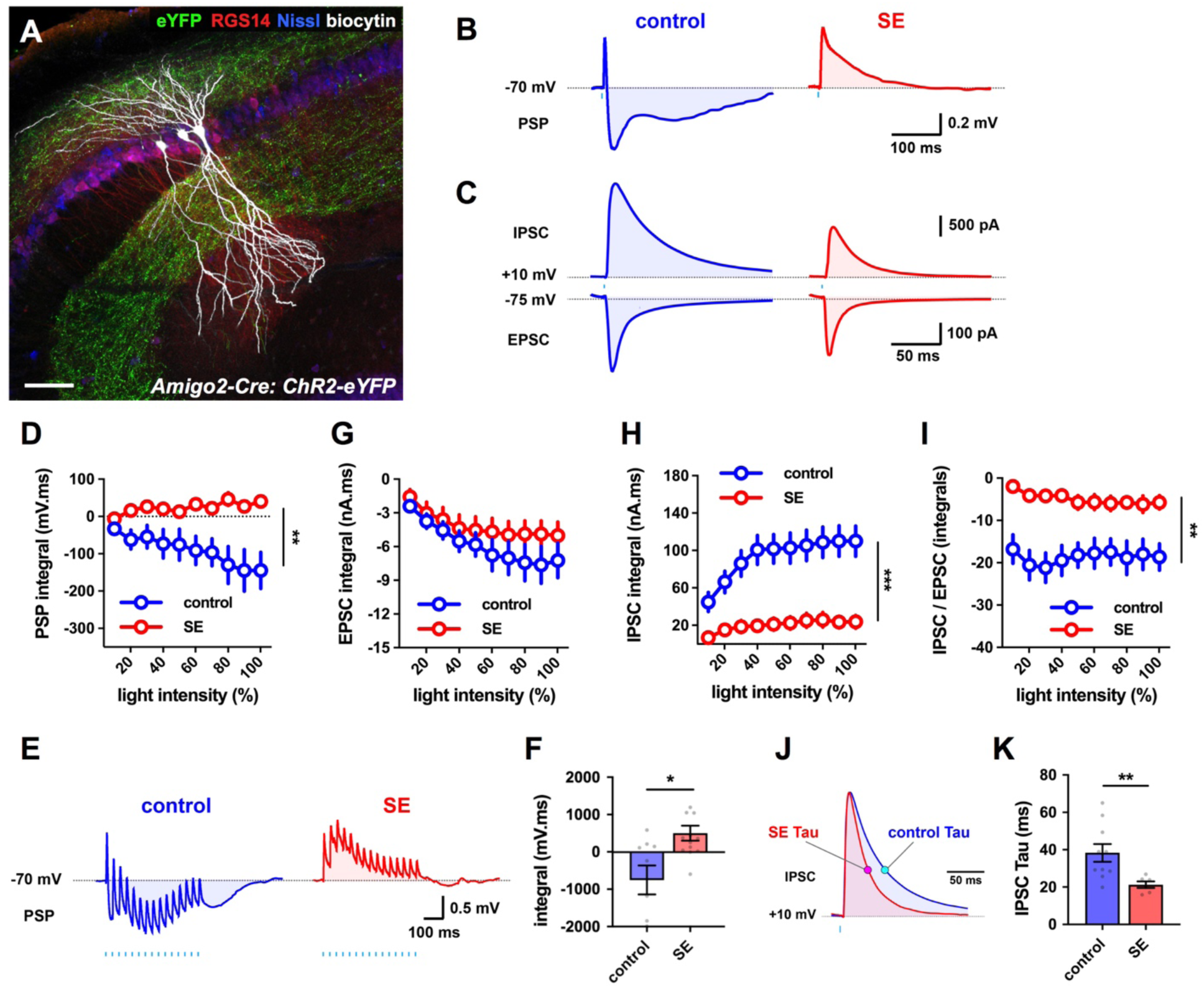
The inhibitory-excitatory balance of the CA2 → CA2 recurrent circuit was reduced in slices from SE. **(A)** Biocytin-filled CA2 PCs (white) in a slice from the intermediate hippocampus, with ChR2-eYFP-expressing CA2 PC axons (green) visible in SO and SR. CA2 PCs were labeled for RGS14 (red) and neuronal somata were visualized with a Nissl stain (blue). Scale bar is 80 μm. **(B)** Representative averaged light-evoked PSPs from CA2 PCs from control and SE mice. **(C)** Representative averaged light-evoked EPSCs and IPSCs from control and SE CA2 PCs. **(D)** The integral of the light-evoked PSP was significantly more positive in CA2 PCs from SE mice. **(E)** Representative averaged PSPs evoked by 15 pulses of light delivered at 30 Hz in cells from control and SE mice. **(F)** The integral of the train-evoked PSP is significantly more positive in cells from SE mice. **(G - I)** Input-output curves of the integral of the light-evoked EPSC (G), the integral of the light-evoked IPSC (H), and the ratio of the integrals of the light-evoked IPSC and EPSC (I). **(J)** Representative averaged light-evoked IPSCs from CA2 PCs with the time constant tau indicated with magenta and cyan markers on the control and SE currents, respectively. **(K)** The time constant tau of the light-evoked IPSC was significantly shorter in cells from SE mice.

In CA2 PCs from control slices, optogenetic stimulation of CA2 recurrent collaterals evoked a triphasic PSP, with an initial EPSP followed by discernable fast and slow IPSPs (figure 3B). PILO-SE treatment produced a large reduction in the inhibitory component of the PSP (figure 3B), similar to the effect on the SR-evoked PSP. Measurement of the integral of the light-evoked PSPs revealed a net hyperpolarization in cells from control mice that was transformed into a net depolarization in Cells from SE mice (figure 3D; mixed-effects model; \*\**P* = 0.0044; n = 15 control, 16 SE). We next applied a train of 15 photostimulation pulses at 30 Hz. In control CA2 PCs the stimulus train evoked PSPs that were dominated by inhibition, resulting in a net hyperpolarization (figure 3E, F). In contrast, photostimulation trains often produced a net depolarization of CA2 PCs from SE mice (figure 3F; \**P* = 0.0208; n = 9 control, 13 SE), with PSPs whose summation exceeded the initial depolarizing response to the first light pulse (one sample t test; \**P* = 0.0362; 13 SE).

Voltage-clamp recordings confirmed the current clamp results (figure 3C, G – K), showing that PILO-SE greatly reduced the IPSC integral (figure 3H; mixed-effects model; \*\*\**P* = 0.0005; n = 17 control, 11 SE), with no change in EPSC integral (figure 3G; mixed-effects model; *P* = 0.3130; n = 16 control, 11 SE). This led to a large decrease in the ratio of the IPSC/EPSC integrals (figure 3I; mixed-effects model; \*\**P* = 0.0023; n = 15 control, 9 SE). PILO-SE also caused a profound decrease in the amplitude of light-evoked IPSCs (mixed-effects model; \*\**P* = 0.0052; n = 17 control, 11 SE) and their duration (figure 3J, K; Mann-Whitney; \*\**P* = 0.0045; n = 10 controls, 6 cells from SE mice). In contrast, PILO-SE had no effect on EPSC amplitude (mixed-effects model; *P* = 0.7769; n = 16 control, 11 SE) or duration (Mann-Whitney; *P* = 0.4378; n = 13 control, 7 SE). Thus, PILO-SE caused a selective loss of inhibition in the CA2 recurrent network, similar to that seen with the CA3 inputs to CA2.

### PILO-SE increases synaptic excitation of CA2 PCs by their mossy fiber inputs from DG

Evidence from clinical and animal studies implicate the dentate gyrus (DG) as a critical network node in temporal lobe epilepsy (Scharfman, 2019). Although the mossy fibers normally provide relatively weak direct excitatory input to CA2 PCs (Kohara et al., 2014; Sun et al., 2017), sprouting of the mossy fiber axons in the CA2 region has been reported both in clinical TLE with MTS (Freiman et al., 2021) and in a mouse model with unilateral MTS-like damage (Häussler et al., 2016). As the functional effects of this sprouting have not yet been determined, we used an optogenetic approach to examine whether PILO-SE alters the mossy fiber input to CA2 PCs. We expressed ChR2 in DG granule cells by crossing the proopiomelanocortin (POMC)-Cre mouse line (McHugh et al., 2007) with a Cre-dependent ChR2-eYFP reporter line (Madisen et al., 2012). ChR2-eYFP+ mossy fiber axons colocalized with the proximal apical dendrites of CA2 PCs which were distinguished from CA3 by their lack of thorny excrescences (figure 4A-C).

**Figure 4.**
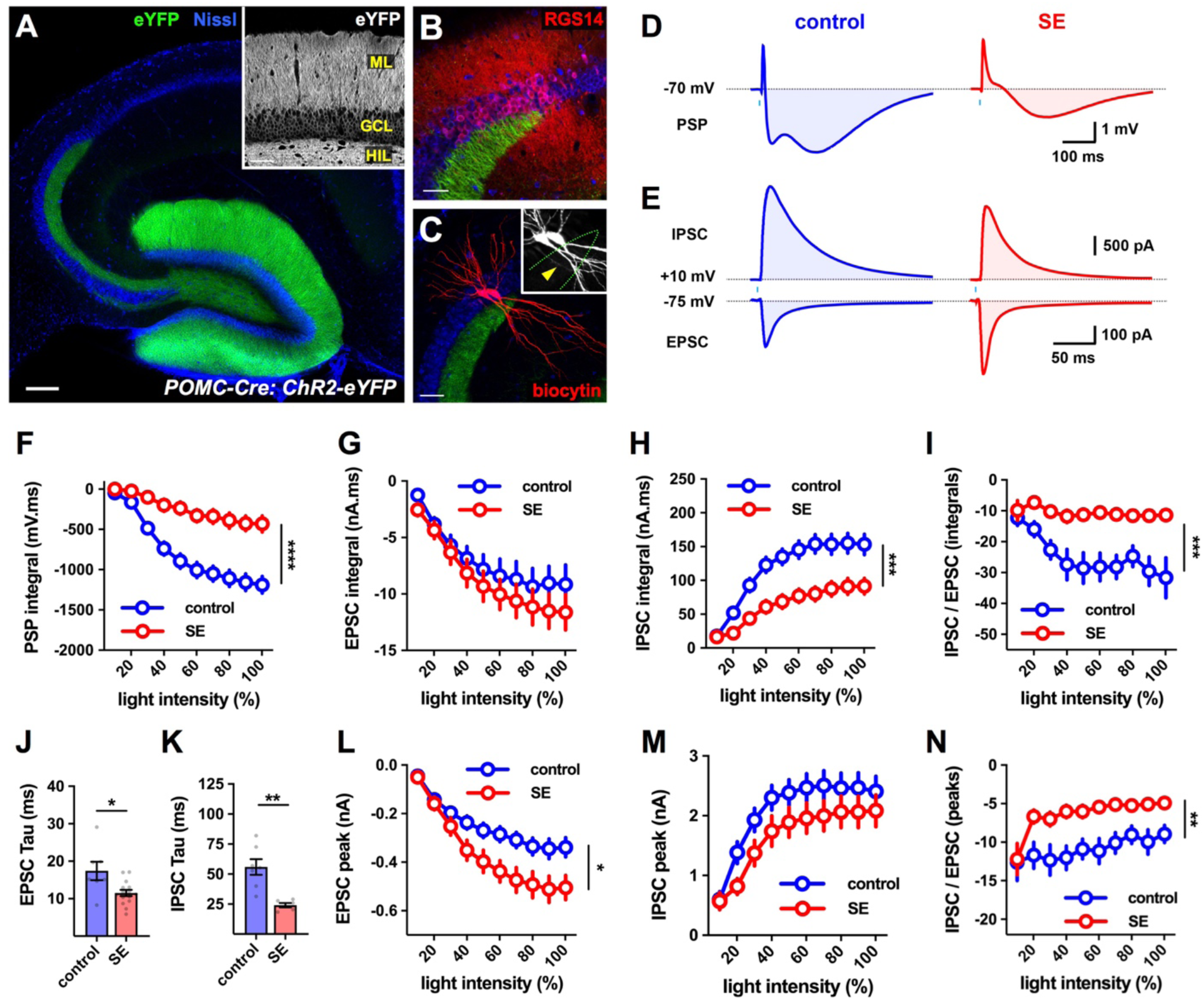
The inhibitory-excitatory balance of the dentate gyrus granule cell mossy fiber pathway to CA2 was reduced after PILO-SE. **(A)** A representative section from a POMC-Cre mouse expressing ChR2-eYFP (green) in DG granule cells, with neuronal somata stained for Nissl substance (blue). Scale bar is 200 μm. Inset, ChR2-eYFP expression in the granule cell layer (GCL), the apical dendrites in the molecular layer (ML), and the mossy fiber axons in the hilus (HIL). Scale bar is 60 μm. **(B)** The CA2 region (defined by RSG14, red) is situated at the end of the mossy fiber projection (Dudek, Alexander, & Farris, 2016). The scale bar is 60 μm. **(C)** Although CA2 PCs (two filled with biocytin, red) lack the thorny excrescences characteristic of CA3 PC proximal apical dendrites (inset, arrowhead), they receive synaptic input directly from GC mossy fibers (defined by past reports and our <4 ms latencies to onset; see (Kohara et al., 2014; Sun et al., 2017). Scale bar is 80 μm. **(D)** Representative averaged light-evoked PSPs in CA2 PCs from control and SE mice. **(E)** Representative averaged light-evoked EPSCs and IPSCs in cells from control and SE mice. **(F)** The input-output curve of the integral of the light-evoked PSP in CA2 from control and SE mice. **(G - I)** Input-output curves of the integral of the light-evoked EPSC (G), the integral of the light-evoked IPSC (H), and the ratio between the integrals of the light-evoked IPSC and the EPSC (I). **(J, K)** The time constants (tau) of the light-evoked EPSC and IPSC were shorter in cells from SE mice. **(L - N)** Input-output curves of the peak amplitude of the light-evoked EPSC in CA2 PCs (L), the peak amplitude of the light-evoked IPSC (M), and the ratio of the peak amplitudes of the light-evoked IPSC and EPSC (N).

In current clamp recordings from CA2 PCs in slices from control mice, photostimulation of mossy fiber axons evoked a triphasic PSP reminiscent of that produced by electrical stimulation in the SR (figure 4D, left). In CA2 PCs from SE mice, the hyperpolarizing phase of the light-evoked PSP was reduced (figure 4D, right) and the integral of the light-evoked PSP was significantly less negative as compared to results from controls (figure 4F; mixed-effects model; control vs SE, \*\*\*\**P* < 0.0001; n = 22 control, 26 SE).

In contrast to the lack of change in excitatory synaptic transmission at the CA3 Schaffer collateral or recurrent CA2 inputs to CA2 PCs, we saw a significant increase in the amplitude of the light-evoked mossy fiber EPSCs in CA2 PCs from SE mice (figure 4E, L; mixed-effects model; \**P* = 0.0334; n = 23 control, 29 SE). Although the peak amplitude of the light-evoked IPSC was not significantly reduced (figure 4M; mixed-effects model; *P* = 0.1457; n = 23 control, 29 SE), both IPSCs and EPSCs exhibited a significantly faster time course of decay in CA2 PCs from SE mice (figure 4J, K; EPSC tau: \**P* = 0.0172, n = 7 control cells and 15 cells from SE mice ; IPSC tau: \*\**P* = 0.0012, n = 7 controls, 6 cells from SE mice). Because of the increased EPSC amplitude, the ratio of the peak IPSC to peak EPSC amplitude was significantly reduced in the epileptic mice (figure 4N; mixed-effects model; \*\**P* = 0.0029; n = 21 control, 28 SE). In addition, because of the increased rate of decay for both the light-evoked IPSC and EPSC, the IPSC integral was much smaller in Cells from SE mice (figure 4H; mixed-effects model; \*\*\**P* = 0.0002; n = 23 control, 29 SE) and the EPSC integral was unchanged (figure 4G; mixed effects model; *P* = 0.3705; n = 23 control, 29 SE), resulting in a net decrease in the ratio of IPSC/EPSC integrals (figure 4I; mixed-effects model; \*\*\**P* =0.0002; n = 21 control, 28 SE). These data suggest that the net excitatory drive of the mossy fiber input to CA2 is enhanced in epileptic mice.

We additionally delivered a 30 Hz photostimulation train (500 ms in duration) and found that in control CA2 PCs light-evoked EPSCs exhibited a small but significant short-term depression (supp. figure 3A, B; the response amplitude at the end of the train (EPSC_15_) was reduced to 55.81 ± 8.494% of the initial EPSC (EPSC_1_); n = 14 control). This EPSC rapidly recovered following the end of the train, returning to its initial level after 500 ms (supp. figure 3A, B). In CA2 cells from SE animals, we observed a dramatic enhancement of short-term depression over the course of the train (supp. figure 3A, B; the amplitude of EPSC_15_ was reduced to 11.85 ± 2.158% of EPSC_1_; control vs SE, two-way ANOVA; \*\*\*\**P* < 0.0001; n = 14 control, 17 SE). Additionally, this depression was longer lasting than observed in cells from control mice as the EPSC was not able to fully recover after a 500 ms interval (supp. figure 3A, B; recovery EPSC amplitude was 109.9 ± 9.45% of EPSC_1_ in controls, and in cells from SE mice the normalized recovery EPSC amplitude was 42.62 ± 5.062% ; n = 14 control, 17 SE).

### PILO-SE strengthens CA2 excitation of CA1

The above results suggest that TLE increases both the intrinsic excitability and net synaptic excitation of CA2 PCs by their mossy fiber, Schaffer collateral, and recurrent excitatory inputs. We next asked whether there was a change in the excitatory synaptic drive from CA2 onto its downstream CA1 PC targets. To address this question, we expressed ChR2 selectively in dorsal CA2 PCs using targeted injections of AAV-DIO-ChR2-eYFP into the CA2 region of Amigo2-Cre mice (supp. figure 2). We then recorded the synaptic response in CA1 PCs to optogenetic stimulation.

We first recorded from deep-layer dorsal CA1 PCs (near the border with stratum oriens, SO) (figure 5A - C), which normally receive stronger CA2 PC input than superficial-layer CA1 PCs (near the border with SR; (Kohara et al., 2014; Valero et al., 2015). We further focused on the CA1c subfield (also termed proximal CA1), near the CA2 border, as this region was more resistant to neurodegeneration that other CA1 subfields in the SE mice.

**Figure 5.**
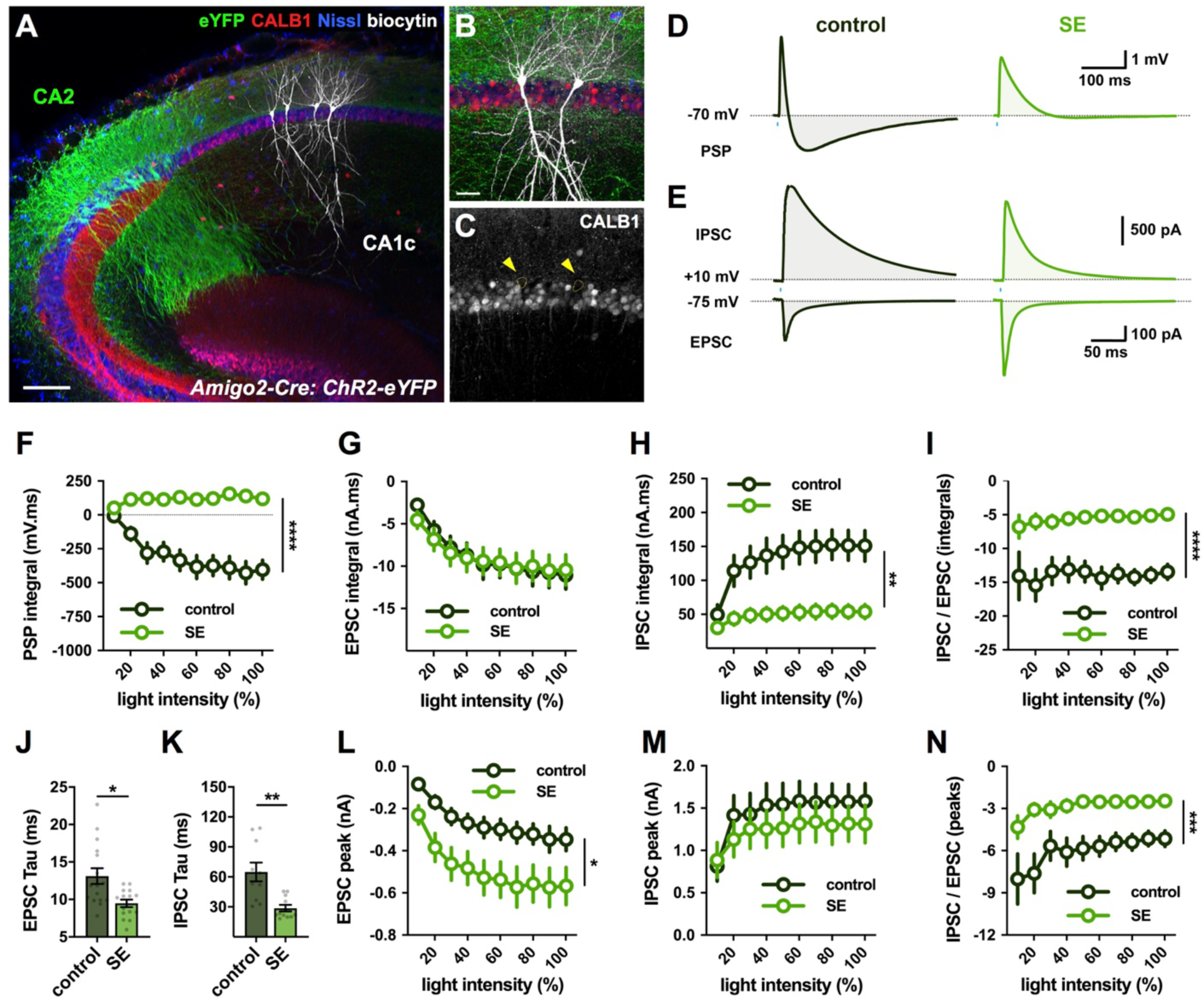
PILO-SE strengthened CA2 excitation of CA1. **(A - C)** A hippocampal slice with biocytin-filled CA1c PCs (white) located in the deep sublayer of SP, adjacent to SO. ChR2-eYFP-expressing CA2 PC dendrites and axonal projections (green) were visible throughout SR and SO. Neuronal somata were labeled with a Nissl stain (blue), and the PC sublayers in CA1 were distinguished using a stain for Calbindin-1 (CALB1, red), which is expressed in CA1_superficial_ cells (Lee et al., 2014). Scale bar is 150 μm in A and 40 μm in B. **(D)** Representative averaged light-evoked PSPs from CA1c_deep_ PCs in slices from control (left, dark green) and SE (right, bright green) mice. **(E)** Representative averaged light-evoked EPSCs and IPSCs from control and SE CA1c_deep_ PCs. **(F)** The integral of the light-evoked PSP was significantly more positive in CA1c_deep_ PCs from SE mice. **(G - I)** Input-output curves of the integral of the light-evoked EPSC, the integral of the IPSC, and the ratio of the IPSC and EPSC integrals. **(J, K)** The time constants (Tau) of the light-evoked EPSC and IPSC were significantly shorter in cells from SE mice. **(L - N)** Input-output curves of the light-evoked EPSC amplitude (L), IPSC amplitude (M), and the ratio of the IPSC and EPSC peak amplitudes (N).

Current clamp recordings from CA1 PCs of control mice revealed a biphasic PSP in response to optogenetic stimulation of CA2 PCs, with an initial brief depolarization followed by a large hyperpolarization, resulting in a net negative PSP integral (figure 5D, left). PILO-SE treatment greatly reduced the hyperpolarization, producing a net positive PSP integral (figure 5D, F; mixed-effects model; \*\*\*\**P* < 0.0001; n = 19 control, 24 SE). Moreover, whereas a brief train of light pulses evoked summating hyperpolarizations and often produced a net negative train integral in cells from control mice, the stimuli elicited a net depolarization and positive integral in PCs from epileptic animals. (figure 6A, B; \*\*\*\**P* < 0.0001; n = 16 control, 23 SE).

**Figure 6.**
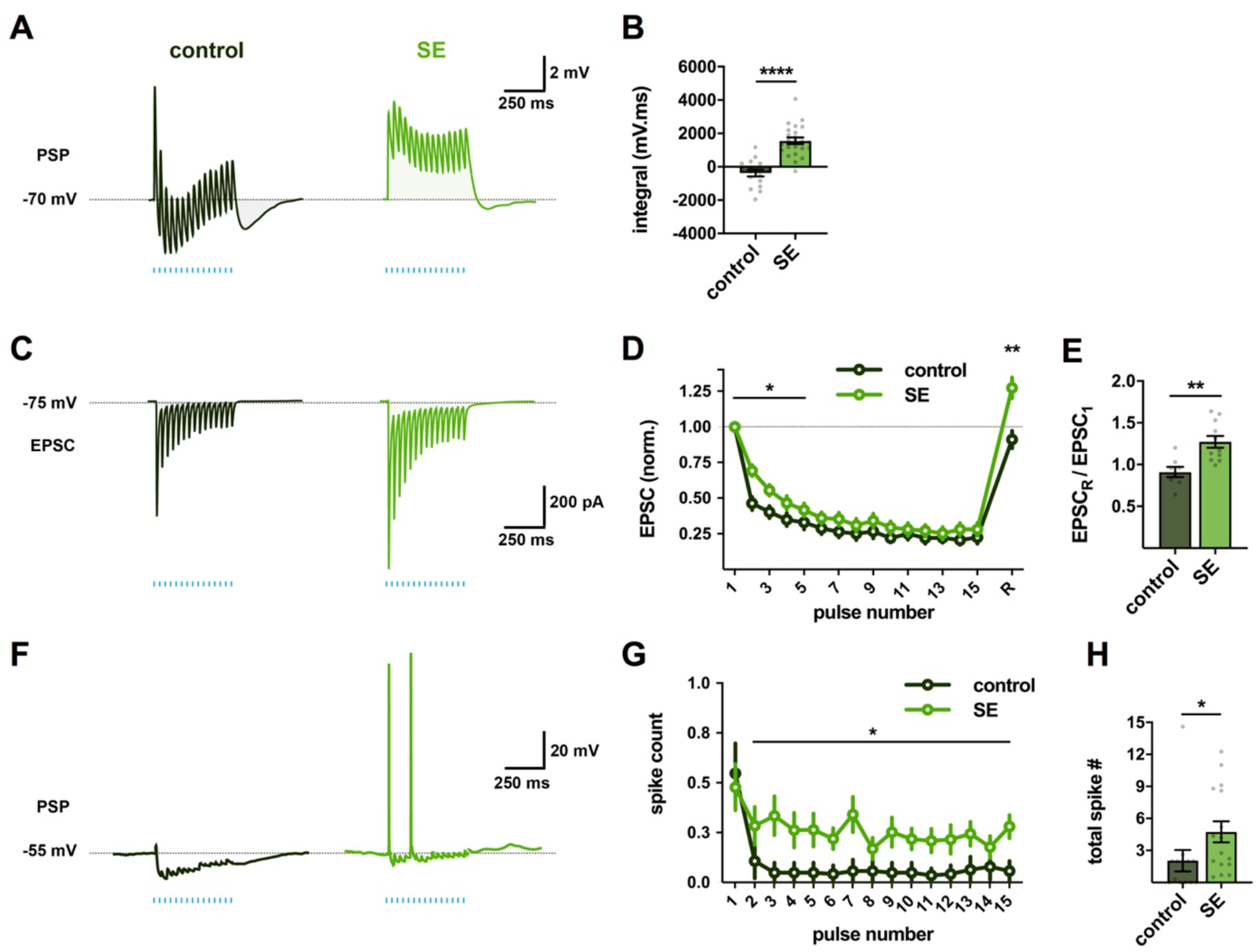
PILO-SE enhanced the ability of CA2 PCs to drive action potential output from CA1 PCs. **(A)** Representative averaged PSPs from control and SE mice evoked by 15 lights pulses delivered at a frequency of 30 Hz for 500 ms. **(B)** The integral of the train-evoked PSP was significantly larger in CA1c_deep_ PCs from SE mice. **(C)** Representative averaged EPSCs evoked by 15 light pulses delivered at a frequency of 30 Hz (blue lines) across 500 ms, recorded from control (dark green) and SE (bright green) CA1c_deep_ PCs. **(D)** Normalized EPSC amplitudes evoked by a train of photostimulation, with 15 pulses delivered at 30 Hz, followed after 500 ms by a single recovery pulse. **(E)** The normalized amplitude of the recovery pulse EPSC was larger in cells from SE mice than in controls. **(F)** Representative photostimulation train-evoked PSPs from CA1c_deep_ PCs held at an initial potential of -55 mV. **(G)** The mean number of action potentials evoked following the first pulse of the photostimulation train was increased in CA1c_deep_ PCs from SE mice. **(H)** The mean total number of action potentials evoked by the 30 Hz photostimulation protocol was significantly increased in CA1c_deep_ PCs from SE mice.

To examine potential changes in the underlying synaptic currents, we next performed voltage-clamp recordings during optogenetic activation. In control animals, photostimulation of CA2 axons evoked a strong monosynaptic EPSC and a long-lasting IPSC in CA1 PCs (figure 5E, left). PILO-SE enhanced significantly the amplitude of the EPSC (figure 5L; two-way ANOVA; \**P* = 0.0184; n = 22 control, 23 SE) but did not alter the amplitude of the IPSC (figure 5M). As a result, the ratio of the IPSC peak currents to EPSC peak currents was reduced (smaller IPSC:EPSC ratio) in CA1c_deep_ PCs from SE mice than from control mice (figure 5N; mixed-effects model; \*\*\**P* = 0.0008; n = 20 control, 20 SE). We also observed that PILO-SE caused a significant faster decay of both EPSCs (figure 5J; \**P* = 0.0147; n = 16 cells from control mice, 15 cells from SE mice) and IPSCs (figure 5K; \*\**P* = 0.0011; n = 10 cells from control mice, 11 cells from SE mice).

Consistent with the decrease in IPSC duration (figure 5K), the integral of the light-evoked IPSC was significantly reduced in PCs from SE mice compared to controls (figure 5H; mixed-effects model; \*\**P* = 0.0025; n = 20 control, 20 SE). In contrast, EPSC integral was unchanged by PILO-SE (figure 5G; two-way ANOVA; *P* = 0.9400; n = 22 control, 23 SE), likely because of offsetting effects of PILO-SE to increase the peak EPSC amplitude while speeding its time course of decay. Consequently, the ratio of the IPSC/EPSC integral was greatly reduced in cells from epileptic mice (figure 5I; mixed-effects model; \*\*\*\**P* < 0.0001; n = 20 control, 20 SE).

PILO-SE also produced a marked decrease in synaptic depression of the EPSC normally observed during a brief train of synaptic stimuli (15 pulses at 30 Hz). with a significant decrease in paired-pulse depression of the first two EPSCs of the train (\*\**P* = 0.0018; n = 8 control, 11 SE). This effect further enhanced the ability of CA2 PCs to excite CA1 PCs, relative to that in control animals (figure 6C, D; two-way ANOVA; first 5 EPSCs; \**P* = 0.0259; n = 8 control, 11 SE), Furthermore, a test stimulus to CA2 delivered 500 ms after the train revealed a short-term potentiation of the EPSC in CA1 PCs from the SE but not control mice (figure 6D, E; \*\**P* = 0.0012; n = 8 control, 11 SE).

In addition to the altered synaptic input, CA1 PCs from SE mice, like CA2 PCs, fired action potentials at a higher rate than controls upon direct current injection (supp. figure 4A; mixed-effects model; \*\*\*\**P* <0.0001; n = 32 control, 46 SE). To determine how altered CA1 excitability and synaptic input from CA2 impacted the ability of CA2 inputs to drive CA1 PC action potential output, we delivered a train of optogenetic stimulation with the CA1 PCs held under current clamp at a membrane potential near threshold (−55 mV), similar to the *in vivo* membrane potentials observed in hippocampal neurons (figure 6F). In slices from control mice activation of CA2 axons with a train of photostimulation evoked only sparse firing in CA1 PCs, with postsynaptic responses dominated by inhibition (figure 6F, G). In contrast, optogenetic stimulation was more effective in exciting CA1c_deep_ PCs from SE mice, with an increase in spike probability following the first stimulus of the train (figure 6G; two-way ANOVA; \**P* = 0.0232; n = 14 control, 16 SE) and an increase in the total number of spikes elicited per train (figure 6H; \**P* = 0.0133; n = 14 control, 16 SE).

In addition to exciting CA1 PCs, CA2 PCs also send an excitatory output to CA3 PCs (supp. figure 2C, D), although this is normally dominated by strong inhibition of CA3 PCs (Kohara et al., 2014; Boehringer et al., 2017). In contrast to the increased net synaptic excitation of CA1 PCs by their CA2 inputs in epileptic mice, PILO-SE caused a decrease in peak amplitude of both the EPSCs and IPSCs recorded from CA3 PCs in response to optogenetic activation of their CA2 PC inputs, with no change in the ratio of inhibition to excitation (supp. figure 5).

### Chemogenetic inhibition of CA2 reduces the frequency of spontaneous seizures

The above results predict that PILO-SE leads to a large increase in the excitability of the cortico-hippocampal circuit through a synergistic combination of actions. These include a widespread decrease in inhibition, increased excitation of CA2 PCs by their mossy fiber inputs, enhanced firing of both CA2 and CA1 PCs, and an enhanced excitatory drive from CA2 to CA1 PCs, which provide the major output from hippocampus to cortex (supp. figure 6). These results led us to test the hypothesis that CA2 PC silencing would reduce seizures in SE mice.

To address this question we used a chemogenetic approach to selectively suppress CA2 PC activity in Amigo2-Cre mice experiencing spontaneous recurrent seizures and asked if the frequency, severity, or other features of seizures were altered compared to controls (figure 7A, see Methods) (figure 7B). Amigo2-Cre mice were injected in CA2 with a Cre-dependent AAV (AAV-DIO-hM4Di-mCherry) expressing the hM4Di inhibitory DREADD (iDREADD; figure 7C). Continuous video EEG recordings began 4 weeks after PILO-SE treatment and continued 24 hr/day, 7 day/week for at least 4 weeks. One group of mice received the DREADD ligand CNO in their drinking water for 2-3 weeks, followed by 2-3 weeks of water without CNO; in a separate group we reversed the order of treatment (figure 7A). In some mice this schedule was then repeated with similar results. After video EEG recordings, mice were perfusion-fixed and brain tissue examined to determine PILO-SE induced pathology and confirm selective expression of virus in CA2 (figure 7D_1_, D_2_). In some experiments, mice were perfusion-fixed at earlier times to confirm CA2-selective DREADD expression throughout the recordings. As previously reported, we observed a classic pattern of neuropathology in SE mice consistent with human MTS, with substantial neuronal loss in the hilus, CA3 and CA1 (Jain et al., 2018; Botterill et al., 2019) and relative survival of CA2.

**Figure 7.**
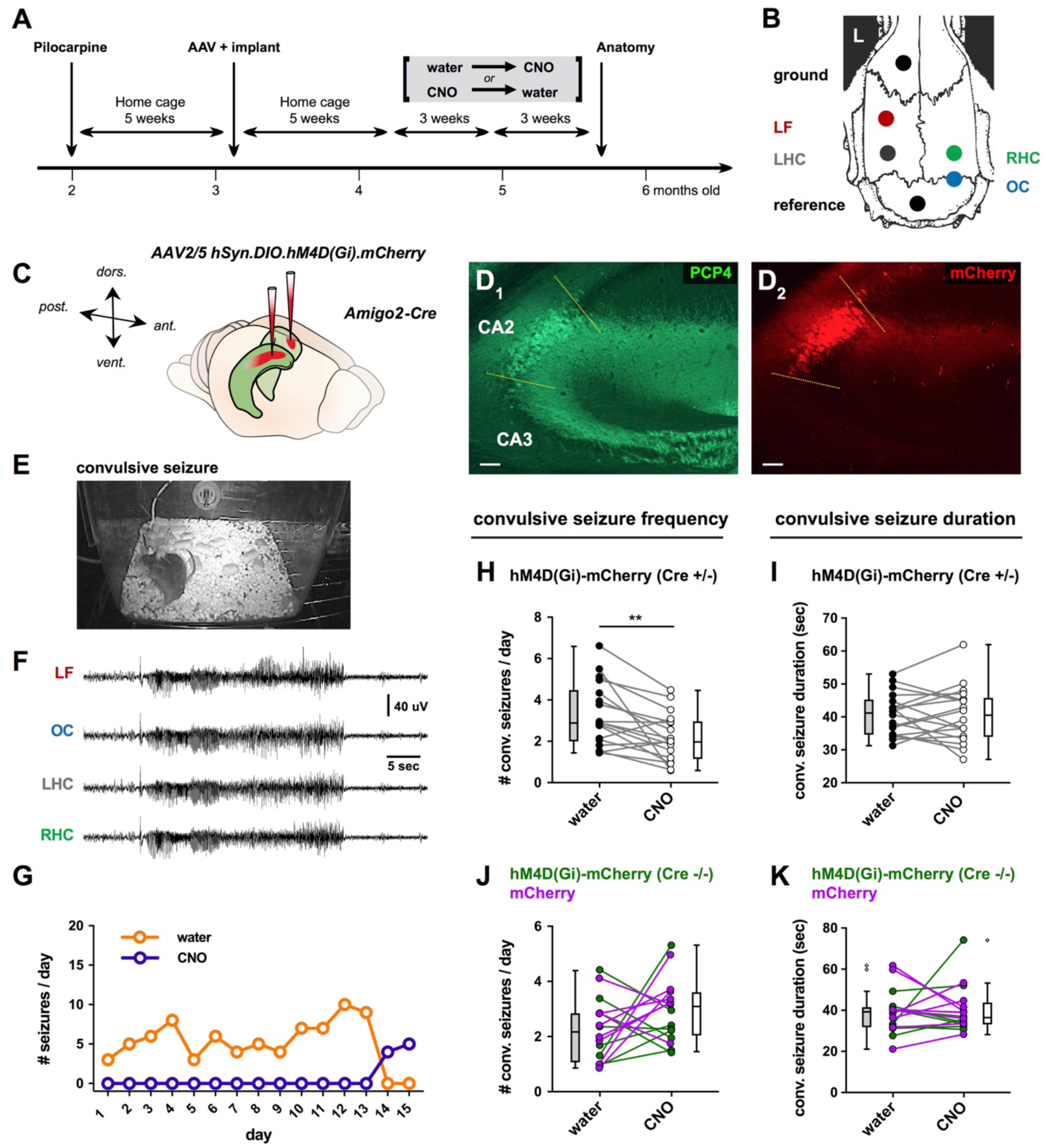
Chemogenetic silencing of CA2 reduced convulsive seizure frequency. **(A)** Timeline of experiments. 5 weeks after PILO-SE Cre-dependent AAV expressing hM4Di-mCherry or mCherry alone were injected in CA2 of Amigo2-Cre mice or wild-type controls. EEG electrodes were implanted at same time. CNO was either present or absent from drinking water for 3-week periods of continuous video EEG recording. Order of CNO delivery was randomized. **(B)** Locations of electrodes for EEG. LF, left frontal; LHC, left hippocampus; RHC, right hippocampus; OC, occipital cortex. **(C)** Locations of bilateral AAV injections in dorsal CA2. **(D)** Representative immunohistochemistry micrographs for CA2 marker PCP4 (D_1_) and AAV-mediated expression of hM4D(Gi)-mCherry (D_2_). **(E, F)** Frame from the continuous video (E) and corresponding EEG (F) during a convulsive seizure in an Amigo2-Cre mouse injected with hM4D(Gi)-mCherry. **(G)** Daily seizure counts in one mouse expressing hM4D(Gi)-mCherry in CA2 during 15 days in the absence of CNO (orange) followed by 15 days with CNO in drinking water (purple). **(H)** Paired analysis of convulsive seizure frequency in the presence of CNO compared with the absence of CNO (water control) in the same Amigo2-Cre mice expressing hM4D(Gi)-mCherry in CA2. CNO reduced convulsive seizure frequency. **(I)** CNO did not alter convulsive seizure duration in Amigo2-Cre mice expressing hM4D(Gi)-mCherry in CA2 (paired t-test; t = 0.0804, df = 17; *P* = 0.9368; n = 18). **(J)** In two groups of control mice (Amigo2-Cre mice expressing mCherry in CA2 (magenta symbols and lines) and wild-type mice (Cre^-/-^) injected with AAV2/5 hSyn.DIO.hM4D(Gi).mCherry) delivery of CNO did not alter convulsive seizure frequency (paired t-test; t = 1.573, df = 15; *P* = 0.1366; n = 16). **(K)** Delivery of CNO did not alter convulsive seizure duration in these control mice (paired t-test; t = 0.3446, df = 15; *P* = 0.7352; n = 16).

PILO-SE treatment reliably induced seizures (9/9 males and 5/9 females; Fisher’s Exact test, p=0.0824), with >90% of seizures associated with stage 4-5 convulsions using the Racine scale (Racine, 1972). EEG activity was similar to seizures in human TLE, with large amplitude, high frequency, rhythmic activity that continued for over 20 sec and was generalized across four electrodes (figure 7E, F). All mice included in the study had frequent convulsive seizures, with total numbers of seizures often exceeding 10 per week (the mean total number of convulsive seizures was 68.94 ± 7.557 over three weeks of recordings, figure 7). In addition, much like human TLE (Baud et al., 2018), seizures could show clustering (supp. fig 7). Most seizures terminated in a prominent postictal depression of the EEG, another characteristic of robust seizures (figure 7F). These characteristics were observed both during periods when CNO was absent from the drinking water and during periods when CNO was delivered. However, the frequency and total number of seizures declined with CNO, as explained below.

Strikingly, periods with CNO treatment showed a marked decrease in convulsive seizure frequency compared with seizure frequency during periods when the drinking water did not contain CNO (exemplified in figure 7G). This decrease was statistically significant when we compared seizure frequency from all mice with and without CNO (figure 7H; paired t-test; t = 3.58, df = 17; \*\**P* = 0.0023; n = 18). The decrease in seizure frequency was not caused by off-target effects of CNO as CNO treatment had no significant effect on seizure frequency in two groups of control mice that did not express iDREADD: Amigo2-Cre^-/-^ mice injected with AAV-DIO-hM4Di-mCherry, and Amigo2-Cre^+/-^ or Cre^-/-^ mice injected with AAV-hSyn-DIO-mCherry (figure 7J). In contrast to the effect on seizure frequency, CNO treatment had no effect on seizure duration, in either iDREADD-expressing PILO-SE mice (figure 7I) or in the two PILO-SE control groups (Figure 7K). Moreover, CNO treatment did not alter seizure severity or the tendency of seizures to occur in clusters (supp. figure 7). Consistent with the decreased convulsive seizure frequency during CNO treatment, total number of convulsive seizures was reduced compared to periods without CNO (paired t-test; **P = 0.0023; n = 18). There was no significant sex difference in either baseline seizure frequency or in the effect of CA2 silencing, so data from male and female mice were pooled. For seizure frequency, a two-way ANOVA with treatment and sex as factors showed an effect of CNO treatment (F(3,48)63.57, p<0.0001), but not sex (F(1,16)0.2305, p=0.6377), and there was no interaction ((F(3,48)1.200, p=0.3197).

In addition to the convulsive seizures, a minority of seizures recorded in the EEG were not associated with behavioral convulsions (figure 8A, B). Similar to its effect on convulsive seizures, CNO treatment of iDREADD-expressing mice also decreased non-convulsive seizure frequency relative to mice that were not treated (figure 8C, E; paired t-test; t = 2.352, df = 16; \**P* = 0.0318; n = 17). There was no effect of CNO on seizure duration (figure 8D), and no effect of CNO in the same control groups discussed above for convulsive seizures (figure 8E, F). As mentioned above, there were few nonconvulsive seizures compared to convulsive seizures (the mean total number of nonconvulsive seizures was 5.00 ± 2.544 over three weeks of recordings). In fact, several mice lacked nonconvulsive seizures entirely (Figure 8C, E).

**Figure 8.**
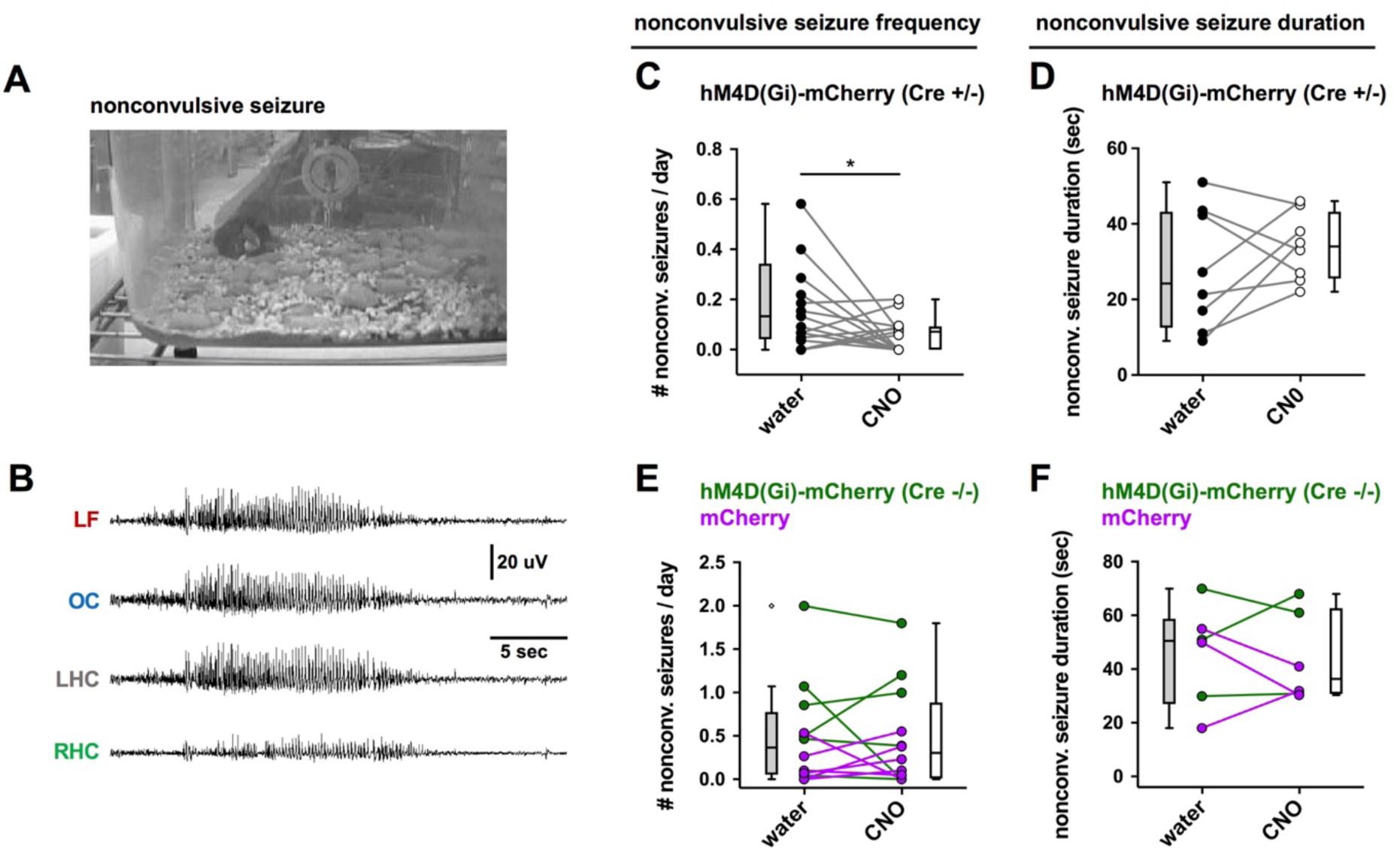
CA2 silencing reduced nonconvulsive seizure frequency. **(A, B)** A representative example of a nonconvulsive seizure in an Amigo2-Cre mouse expressing hM4D(Gi)-mCherry in CA2. Despite the absence of behavioral indicators (A), EEG revealed significant seizure activity in all electrodes (B). **(C)** The presence of CNO reduced nonconvulsive seizure frequency compared to the absence of CNO in these mice. **(D)** The presence of CNO did not alter seizure duration (paired t-test; t = 0.6743, df = 6; *P* = 0.5252; n = 8). **(E)** CNO did not reduce nonconvulsive seizure frequency in the two control groups of mice (described in the figure; paired t-test; t = 0.1333, df = 11; *P* = 0.8963; n = 12). **(F)** CNO did not reduce nonconvulsive seizure duration in the two control groups of mice (paired t-test; t = 0.2927, df = 5; *P* = 0.7815; n = 6).

## DISCUSSION

Here we provide the first direct evidence that CA2 PCs play an important role in seizure generation, based on findings that chemogenetic silencing of CA2 reduces seizure frequency in the PILO-SE model of temporal lobe epilepsy. In principle, this could reflect a relatively passive role of CA2, in which the survival of CA2 PCs with normal excitability provides a pathway for propagation of seizure activity arising from abnormal activity in upstream regions. However, CA2 could also serve a more active role, in which alterations to CA2 intrinsic excitability and/or synaptic circuitry would enhance local seizure generation and/or seizure propagation. The latter is suggested by data from human TLE specimens resected because of intractable epilepsy (Wittner et al., 2009).

Support for an active role for CA2 is also provided by our *ex vivo* hippocampal recordings, which reveal a coordinated increase in CA2 firing and enhanced net excitatory synaptic input to and output from CA2. Thus, our *ex vivo* slice studies showed a general loss of synaptic inhibition both within CA2 and other hippocampal regions that would enhance propagation of electrical activity through the hippocampal circuit. At the same time we observed a significant increase in the direct excitatory synaptic input that CA2 receives from the dentate gyrus (DG) mossy fibers. While not considered part of the classic trisynaptic pathway, the DG → CA2 circuit can nevertheless support the propagation of activity through the hippocampal network . In normal brain function, the mossy fiber pathway directly excites CA3 PCs but provides relatively weak excitatory input to CA2 in a parallel trisynaptic circuit (Kohara et al., 2014; Sun et al., 2017). DG granule cells are highly resistant to the cell death characteristic of mesial temporal sclerosis (MTS), and as such the DG has long been thought to function as a critical network node in TLE that gates excitatory input from the entorhinal cortex to downstream hippocampal neuronal populations (Krook-Magnuson et al., 2015; Scharfman, 2019). However, it has been unclear how DG may participate in seizures since its major synaptic target, CA3 PCs, undergoes marked degeneration in TLE accompanied by MTS (Blümcke et al., 2013; Steve et al., 2014; Winawer et al., 2007).

Our finding of enhanced mossy fiber excitation of CA2, together with increased CA2 firing and decreased inhibition, would increase the efficacy of the DG in triggering CA2 spiking, effectively bypassing damaged CA3 tissue. Moreover, our finding of an increased excitatory synaptic drive from CA2 PCs to CA1 PCs, accompanied by a decrease in inhibition, would amplify the ability of CA2 to excite CA1. Together our results suggest a powerful role of CA2 as a proconvulsant subfield in TLE.

### Chemogenetic modulation of CA2 PCs decreased the frequency of spontaneous seizures in the PILO-SE model

A proconvulsant role of CA2 due to its altered excitability and enhanced net excitatory synaptic input and output provides a potential explanation for how CA2 activity contributes to seizure activity *in vivo*. Notably, we found that selective and chronic chemogenetic silencing of CA2 PCs led to a significant decrease in spontaneous convulsive and nonconvulsive seizure frequency. However, CA2 inhibition affected neither seizure duration nor seizure severity. Therefore CA2 may exert the greatest influence over seizures that begin in hippocampus, and thus CA2 modulation may be less effective when seizures emerge from extrahippocampal regions. Indeed, data from both the PILO-SE model and clinical seizure recordings suggest that seizures in TLE are generated from several areas, not only the hippocampus (Ogren et al., 2009; Spencer, 2002; Wyeth, Nagendran, & Buckmaster, 2020). In our EEG recordings it was not possible to discriminate whether seizures began in hippocampus or elsewhere because the electrodes were subdural, relatively remote from the areas like hippocampus.

### CA2 promotion of seizure activity in epileptic mice reflects its function in the healthy hippocampus

A strong control by CA2 of pathological hippocampal network activity such as temporal lobe seizures is consistent with recent investigations into the role of CA2 in healthy animals (Kay & Frank, 2019; Lehr et al., 2021). Thus, CA2 PCs play an important role in generating sharp wave ripples (SWRs), brief periods of synchronous firing observed during slow wave sleep. Although SWRs were presumed to originate primarily in the CA3 recurrent network, a significant fraction of SWRs arise in CA2, with an increase in CA2 firing rates often preceding the increase in CA3 during the buildup to a SWR (Oliva et al., 2016). CA2 PCs, like those in CA3, form recurrent excitatory connections (Okamoto & Ikegaya, 2018). Although acute optogenetic inhibition of CA2 decreases the frequency of SWR occurrence (Oliva et al., 2020), more prolonged inhibition of CA2 using chemogenetics can lead to a paradoxical increase in SWR occurrence (Alexander et al., 2018), consistent with findings that CA2 backprojections recruit powerful inhibition of CA3 PCs (Boehringer et al., 2017; Kohara et al., 2014). Additionally, permanent tetanus toxin-mediated CA2 silencing may produce compensatory changes that enhance the excitability of the CA3 recurrent network through reduced inhibition (Boehringer et al., 2017). We speculate that in TLE, when synaptic inhibition is compromised, CA2 cannot constrain CA3 excitability and the CA2 and CA3 subfields together act as a hyperexcitable hub. There is also a large literature suggesting that the DG generates seizures in the PILO-SE model (reviewed in Scharfman, 2019), and in cases where the CA3 subfield exhibits neurodegeneration, such seizures would depend on passage to CA2 to propagate from the hippocampus and develop into a prolonged (>20 sec) generalized seizure. Importantly, in resected human TLE tissue the resistant CA2 subfield generates epileptiform activity in parallel to, and independently from, the subiculum (Wittner et al., 2009).

### Altered intrinsic excitability of CA2 PCs in TLE

Direct somatic current injection caused CA2 PCs from epileptic mice to fire at a higher rate than cells from control mice (figure 1B, C), as previously reported for CA1 and CA3 PCs (Arnold, Mcmurray, Gray, & Johnston, 2019; Barmashenko, Hefft, Aertsen, Kirschstein, & Köhling, 2011). The input resistance (R_in_) was increased and the membrane capacitance (Cm) was decreased in CA2 PCs from epileptic mice (figure 1E, F). As a result, synaptic and experimentally-applied currents may cause a greater and more rapid change in membrane potential. Indeed, though the action potential voltage threshold was not altered, the rheobase current was reduced (figure 1L). However, these altered membrane properties may enhance both depolarizations and hyperpolarizations and the net effect on CA2 PC activity *in vivo* is unclear.

Although the action potential voltage threshold was unchanged in SE mice, indicating a lack of direct effect on fast voltage-gated Na^+^ channels, the increase in the fast afterhyperpolarization (AHP) that follows a single action potential (figure 1M) may facilitate repetitive high-frequency firing by increasing the rate of sodium channel recovery from inactivation (Jaffe & Brenner, 2018). We also found an increase in the amplitude of the hyperpolarization-activated voltage sag (figure 1I, J), suggesting an enhancement of currents mediated by hyperpolarization-activated cyclic nucleotide-gated (HCN) channels in CA2 PCs following SE (Srinivas et al., 2017). This stands in sharp contrast to numerous reports of HCN downregulation in CA1 PCs from epileptic rodents (Arnold et al., 2019; Wolfart & Laker, 2015), though upregulation of HCN currents has been observed in DG granule cells in surgically resected human TLE tissue (Stegen et al., 2012). Thus, examination of CA2 PC intrinsic properties revealed a wide array of alterations in epileptic mice (figure 1, supp. table 1), many of which are expected to increase the excitability of CA2 PCs *in vivo*.

### Increased excitability of synaptic inputs to and output from CA2 PCs in epileptic mice

CA2 normally receives strong direct excitatory input from layer II neurons of entorhinal cortex (EC LII), weak direct excitatory input from the mossy fibers, and strong excitatory input from CA3 that is curtailed by strong inhibition (Chevaleyre & Siegelbaum, 2010; Nasrallah et al., 2019; Sun et al., 2017). In PILO-SE, we saw heterogeneous effects on the strength of these inputs, with no change in the EPSC or IPSC evoked by entorhinal cortical inputs, an increased EPSC and decreased IPSC in response to mossy fiber excitation, and a decreased IPSC with unchanged EPSC in response to activation of the CA3 Schaffer collaterals. The more rapid decay of IPSCs evoked by Schaffer collateral stimulation and mossy fiber activation in CA2, or by activation of CA2 axons in CA1, may reflect decreases in inhibitory current mediated by GABA_A_ receptors, as GABA_B_-mediated inhibition is observed at a longer latency after stimulation (for example see figure 2B, C). Precedent for a loss of inhibition in CA2 may be found in a prior investigation in which whole-cell recordings from CA2 PCs were performed in human hippocampal tissue surgically resected either from patients with tumor-associated TLE or from patients with TLE with mesial temporal sclerosis (MTS): intriguingly, upon stimulation of the Schaffer collaterals prominent feedforward inhibitory postsynaptic potentials were observed in CA2 PCs from tumor-associated TLE tissue, but not in CA2 PCs from TLE MTS tissue (Williamson & Spencer, 1994).

In addition, at the DG → CA2 circuit we observed a substantial increase in short-term depression that could reflect an elevated presynaptic release probability (supp. figure 3), as reported previously at synapses formed by sprouted mossy fibers (Hendricks, Chen, Bensen, Westbrook, & Schnell, 2017; Hendricks, Westbrook, & Schnell, 2019). At the CA2 → CA1c_deep_ circuit we found short-term synaptic potentiation not seen in control tissue (figure 6C - E). These data suggest that in epileptic mice, CA2 PNs are intrinsically hyperexcitable and poised to convey excitatory activity along the alternate trisynaptic circuit from DG → CA2 → CA1. Since CA3 neurons are the primary recipient of mossy fiber input and the major contributor to the Schaffer collateral pathway, enhancement of the DG → CA2 → CA1 parallel pathway may represent a compensatory response to the depletion of CA3 neurons. The loss of mossy fiber targets in CA3 may trigger a compensatory increase in mossy fiber input to CA2. This model is consistent with prior suggestions that, following the loss of normal targets in the hilus and CA3, granule cell mossy fibers sprout and establish reorganized synaptic connections onto surviving target neurons in the dentate gyrus, including other granule cells (Schmeiser, Zentner, Prinz, Brandt, & Freiman, 2017). Similarly, the loss of CA3 Schaffer collateral synapses on CA1 PCs and the decreased CA2 synaptic output to CA3 may cause a compensatory increase in excitatory input from CA2 to CA1. These data together suggest that in epileptic mice there is a rebalancing of CA2 output away from the backprojection to CA3 and in favor of stronger activation of CA1. Importantly, CA2 PC axons project along both the transverse and longitudinal axes of the hippocampus (supp. fig 2; see Meira et al., 2018; Okuyama, Kitamura, Roy, Itohara, & Tonegawa, 2016; Tamamaki, Abe, & Nojyo, 1988), and additional target numerous extrahippocampal structures (Cui, Gerfen, & Young, 2013). CA2 projections therefore potentially provide an extensive circuit substrate for the propagation of epileptiform activity.

Our finding that the entorhinal cortical synaptic excitation of CA2 is unchanged in TLE is consistent with findings that there is an increased survival of EC LII neurons relative to those in EC LIII in both human TLE tissue (Fu Du et al., 1993) and numerous animal models of epilepsy (F. Du, Eid, Lothman, Kohler, & Schwarcz, 1995; Janz et al., 2017; Tolner et al., 2007; Wozny et al., 2005). Thus, in some cases of TLE with hippocampal neurodegeneration, not only do CA2 PCs survive but so too do the EC LII neurons that comprise their principal input. Furthermore, CA2 PCs recruited by EC LII directly excite downstream CA1 PCs in a powerful disynaptic circuit (Bartesaghi, Migliore, & Gessi, 2006; Chevaleyre & Siegelbaum, 2010). Although we did not find functional alterations to the EC LII input to CA2, this circuit nevertheless may serve as an important route for seizure propagation through the hippocampal network.

## Conclusion

In summary, our data provide compelling support for the hypothesis that CA2 circuits become pathologically hyperexcitable following PILO-SE as a result of changes in both intrinsic and extrinsic excitability, and this contributes to the emergence of seizures in chronically epileptic mice. The combined effect of these changes, summarized in supplementary figure 6, may facilitate the emergence of seizures in the hippocampus or their propagation through the hippocampal-entorhinal cortical network via CA2-centered pathways. Thus, our findings suggest that the surviving CA2 PC population may function as a resilient network node within the cortico-hippocampal loop that supports seizure generation and propagation, and CA2 may therefore be an important novel target for the treatment of drug-resistant seizures in TLE.

## Acknowledgements

This work was supported by grant R01NS106983 from NIH (PIs, H.E.S. and S.A.S.) and grant F31NS113466(A.C.W.).

## AUTHOR CONTRIBUTIONS

A.C.W., J.J.L., S.J. and P.L. acquired and analyzed data. A.C.W., H.E.S. and S.A.S. designed the experiments and wrote the manuscript.

## METHODS

### I. Animals

All procedures were performed in accordance with the Nathan Kline Research Institute for Psychiatric Research (NKI) and Columbia University Institutional Animal Care and Use Committees (IACUC). Adult male and female mice (8-12 weeks-old) were housed in a temperature and humidity-controlled environment with a 12-hour light/dark cycle with food and water provided *ad libitum*. For in vivo studies, Amigo2-Cre^+/-^ mice from the Siegelbaum laboratory (C57BL/6J background) were initially bred at NKI with wild-type (WT) C57BL/6N mice (stock # 00664; Jackson Laboratory) and backcrossed with the C57BL/6N line for several generations before use. For *ex vivo* studies at Columbia University, the Amigo2-Cre^+/-^ mice from the Siegelbaum laboratory were F1 generation hybrids resulting from a cross between C57BL/6J mice (stock # 000664; Jackson Laboratory) and 129S1/SvlmJ mice (stock # 002448; Jackson Laboratory). Genotyping was performed using tail tip samples sent to the New York University Mouse Genotyping Core facility (NKI) or GeneTyper (Columbia University). In a small cohort at the outset of the study it was confirmed that genotypes of mice were the same using the two genotyping facilities.

### II. Induction of epilepsy

Epileptic mice were generated after pilocarpine-induced SE (PILO-SE) for *in vivo* and *ex vivo* studies. *In vivo* studies used hM4Di (inhibitory DREADDs; iDREADDs) to selectively silence CA2 PCs during chronic seizures. *Ex vivo* studies used hippocampal slices and associated techniques to clarify potential mechanisms underlying the role of CA2 in epileptic conditions. Initial studies using identical mice and methods at NKI and Columbia University led to greater mortality at Columbia University, so Columbia University modified methods to reduce mortality. This led to slightly different methods but each location generated robust epileptic mice. One change that was made was Columbia University used a slightly different background strain to reduce mortality, a F1 generation hybrid resulting from a cross of the C57BL/6J strain (high mortality) and 129S1/SvlmJ mice (lower mortality). However, these mice did require a higher dose of pilocarpine (see below for dose). A higher dose was also likely to be a result of a higher body weight in the mixed background strain.

#### A. PILO-SE induction for *in vivo* experiments

Induction of SE with the convulsant pilocarpine was done in cohorts of 2-4 mice with the experimenter blinded to the experimental group. Baseline vEEG recordings were acquired for at least 1 hr to capture a wide range of EEG signals associated with various behavioral states (e.g., exploration, grooming, and rest). Following the baseline period, mice were injected with the peripheral muscarinic antagonist scopolamine methyl nitrate (1 mg/kg s.c.; #S2250, Sigma Aldrich) to reduce the peripheral effects of pilocarpine. The β2-adrenergic agonist terbutaline hemisulfate (1 mg/kg s.c.; #T2528, Sigma Aldrich) was also administered to support respiration. Ethosuximide (150 mg/kg s.c.; #E7138, Sigma Aldrich) was administered to reduce the occurrence of brainstem seizures which can lead to mortality (Iyengar et al., 2015). PILO-SE was induced by injecting pilocarpine hydrochloride (250 mg/kg s.c., #P6503, Sigma Aldrich). All mice were injected with diazepam (5 mg/kg, s.c.; NDC# 0409-3213-12, Hospira) 2 hrs after the pilocarpine injection to reduce the severity of SE, which appears to prevent morbidity and mortality after SE (Goodkin & Kapur, 2009; Iyengar et al., 2015). Mice were injected with 1 mL (s.c.) of lactated Ringer’s solution (Aspen Veterinary Resources) at this time to support hydration. Our previous studies suggest that SE is most intense for several hours after the pilocarpine injection, but there is continued spiking in the EEG overnight (Iyengar et al., 2015; Jain, LaFrancois, Botterill, Alcantara-Gonzalez, & Scharfman, 2019). For all *in vivo* experiments, stock solutions were freshly prepared in 0.9% NaCl in dH_2_O (saline; i.e., CNO, scopolamine, terbutaline) or phosphate buffered saline (i.e., ethosuximide).

#### B. PILO-SE induction for *ex vivo* experiments

All drugs were administered intraperitoneally (i.p.). Mice were first administered methylatropine bromide (5 mg/kg, i.p.; Millipore Sigma #M1300000) to suppress peripheral cholinergic activation from pilocarpine hydrochloride. Pilocarpine was administered 30 mins later (350 mg/kg, i.p.; Sigma Aldrich #P0472) and mice were closely and continually monitored for behavioral indicators of seizures. The onset of SE typically occurred between 30 and 60 mins following pilocarpine treatment and was defined as a convulsive seizure (stage 3, 4, or 5 on the Racine seizure scale (Racine, 1972)) that lasted continually for at least 5 mins and did not fully subside for several hours. Diazepam (5 mg/kg, i.p.; McKesson #636203) was administered 1 hr after SE onset to curtail seizures. If mice did not exhibit SE following pilocarpine injection (non-SE) they were administered diazepam at a similar delay after pilocarpine as SE mice, 2 hrs after pilocarpine. In all cases diazepam was followed 20 mins later by levetiracetam (100 mg/kg, i.p.; West-Ward NDC #0143-9673-10). Control mice were given an identical course of drug treatment to non-SE mice, except they were not administered pilocarpine. Thus, control animals received methylatropine bromide, then after 150 mins were administered diazepam and at 170 mins were administered levetiracetam. Immediately after levetiracetam SE and non-SE mice were transferred to a heated and humidified veterinary intensive care unit (ThermoCare #FW-1), where they were closely monitored, provided with dietary supplements, and given subcutaneous hydration (lactated Ringer’s solution; ICU Medical NDC #0409-7953-03) until they showed normal locomotion and feeding (typically within 1-3 days). After recovery all mice were kept in standard group housing, except where aggression between cage-mates was observed in which cases aggressors were removed. In the cohort of mice used for *ex vivo* experiments, pilocarpine treatment resulted in three outcomes: acute mortality due to generalized tonic-clonic seizure leading to tonic hindlimb extension and death in 102/367 mice (27.8%), minor isolated seizures but without SE in 110/367 mice (30%), or SE in 155/367 mice (42.2%).

### III. *In viv*o experiments with epileptic mice

#### A. Stereotaxic injection of AAV

Mice were initially anesthetized with 5% isoflurane (Aerrane, Henry Schein). The mice were then immediately secured in a rodent stereotaxic apparatus (Model #502063, World Precision Instruments). A homeothermic blanket system maintained body temperature at 37 °C (Harvard Apparatus). Isoflurane (1-2%) was mixed with oxygen and delivered through a nose cone attached to the stereotaxic apparatus. Buprenex (Buprenorphine, 0.1 mg/kg, s.c.) was delivered prior to any surgical manipulations to reduce discomfort. The scalp of each mouse was then shaved and swabbed with Betadine (Purdue Products). Lubricating gel was applied to the eyes to prevent dehydration (Patterson Veterinary).

After a midline scalp incision, a surgical drill (Model C300, Grobert) was used to make two craniotomies for viral injections. All stereotaxic coordinates for craniotomies are described in anterior-posterior (AP) and medial-lateral (ML) coordinates (in reference to Bregma). Craniotomies were made over dorsal CA2 bilaterally (−1.9 mm AP, ± 1.4 mm ML). This injection site was chosen to maximize specific expression in CA2.

A 33-gauge infusion needle (#C315I-SPC, Plastics One) attached to a 0.5 µl Hamilton syringe was lowered from the skull surface 1.5 mm into each relatively anterior site in the hippocampus to target dorsal CA2. Each site was injected with 150 nL of virus at a rate of 40 nL/min. The needle remained in place for an additional 5 min after each injection to allow for diffusion of the virus and then it was slowly removed.

Next, subdural screw electrodes (0.10” length stainless steel jeweler’s screws #8209, Pinnacle Technology) were secured in the two anterior craniotomies positioned over the left and right dorsal hippocampus. A subdural screw electrode was also secured in the right occipital cortex (−3.5 mm AP, +2.0 ML). In addition, a subdural screw electrode was secured in a craniotomy made over the left frontal cortex (0.0 mm AP, -2.6 mm ML). Last, screw electrodes were secured in craniotomies made over the right olfactory bulb (+2.3 mm AP, +1.8 mm ML) which served as a ground and the cerebellum (−5.7 mm AP, -0.5 mm ML) which served as a reference electrode. The subdural screw electrodes were attached to an 8-pin connector that was centered over the skull and secured with dental cement. Mice were transferred to a clean cage at the end of surgery and placed on a heating blanket (37 °C) until fully ambulatory.

#### B. Video-EEG and CNO treatment

Mice were housed individually in a standard laboratory cage within a room where the video-EEG (vEEG) equipment was housed so they would acclimate to the recording environment. After the recovery period, each mouse was individually placed into a 21 cm x 19 cm transparent cage comparable to a standard laboratory cage. A pre-amplifier was inserted into the 8-pin headcap and connected to a multichannel commutator (Pinnacle Technology). This tethered EEG system allowed for free range of movement throughout the entire recording cage. EEG signals were acquired at 500 Hz and bandpass filtered at 1-100 Hz in Sirenia Acquisition software (Pinnacle Technology). Simultaneous video recordings synchronized with the EEG record were captured using an infrared LED camera (#AP-DCS100W, Apex CCTV) for offline analyses.

CNO (Sigma-Aldrich) was added to the drinking water at a 0.4 mg/ml concentration. For a 20 mg mouse the water delivered approximately 10 mg/kg per day, given an average volume of water consumption of 6 mL per day. The CNO stock solution (1 mg/ml dissolved in water) was made every 3 days and refrigerated between uses. We selected a 10 mg/kg/day dose of CNO because this dose has been widely used in the literature to inhibit diverse cell types throughout the brain with minimal to no off-target effects (Mahler & Aston-Jones, 2018; Smith, Benison, Bercum, Dudek, & Barth, 2018). Importantly, we controlled for potential off-target effects of CNO by also injecting Cre^-/-^ mice with the same dose of CNO as iDREADD-expressing mice (MacLaren et al., 2016). In prior studies we found no evidence of non-specific effects of the 10 mg/kg dose of CNO (Botterill et al., 2019, 2021).

All vEEG recordings were analyzed offline using Sirenia Seizure Pro (v. 1.7.9, Pinnacle Technology). Seizures were defined as rapid and rhythmic (>3 Hz) deflections in all EEG channels that lasted >5 sec (Botterill et al., 2019; Iyengar et al., 2015; Jain et al., 2019) and were at least 3 standard deviations above the baseline root mean square (RMS) amplitude (Iyengar et al., 2015). Seizures were considered convulsive if the video record showed behaviors consistent with stages 3-5 on the Racine scale (stage 3, unilateral forelimb clonus; stage 4, bilateral forelimb clonus with rearing; stage 5, stage 4 followed by loss of posture; (Racine, 1972). Seizures were considered non-convulsive if the EEG criteria were met, but no stage 3-5 behaviors were detected in the video record. Nonconvulsive seizures were occasionally accompanied by small movements but often occurred during sleep or immobility. SE was defined as the first seizure that was severe (large amplitude EEG deflections appearing in all 4 electrodes simultaneously, stage 3-5) with the seizure activity in the EEG persisting continuously for >5 min, and seizure-associated behavior persisting for >5 min, a standard definition (Goodkin & Kapur, 2009; Iyengar et al., 2015). Five min was chosen because if the abnormal activity persistent for 5 min it lasted for several hours. Note that during SE the seizure-associated behavior included up to 10 stage 3-5 convulsions. Between convulsions behavior included twitching and abnormal body movement. Cessation of convulsions was indicated by a return to grooming and feeding although animals showed persistent EEG spiking, seizure activity and convulsions intermittently for the next 24 hours (Botterill et al. 2019 cell rep).

#### C. Anatomical assessments

Following deep anesthesia by isoflurane inhalation and a terminal dose of urethane (2.5 g/kg, i.p.), the abdominal cavity was opened and a 26 g butterfly needle inserted into the ventricle. After cutting the left atria, 0.9% NaCl (30 mL) was perfused into the heart with a peristaltic pump (Gilson) on a high setting (1 mL/min). Fixative (4% paraformaldehyde in 0.1 M Tris buffer, pH 7.6; 30 mL) was perfused immediately thereafter. After storing the body at 4° C for over 30 min, the brain was removed and postfixed in paraformaldehyde for at least 24 hrs.

Sections (50-µm thick) were cut with a vibratome (Leica 1200; Leica Biosystems) in the coronal plane and alternate sections were stained using Cresyl violet (Scharfman, 2002) or with immunohistochemical procedures using markers of CA2 (PCP4, RSG14; described further below). Sections were coverslipped with Permount (Fisher) and viewed with an upright microscope (BX61; Olympus of America) using a digital camera (Infinity3-6URC; Lumenera) and associated software (Infinity; Lumenara).

### IV. *Ex vivo* experiments with epileptic mice

#### A. AAV for optogenetics

Immunohistochemistry and *ex vivo* electrophysiology were performed in the chronic phase of PILO-induced epilepsy, at least 6 weeks after status epilepticus, when mice were 14 – 18 weeks old. Surgeries were performed at least three weeks after pilocarpine treatment. Data from male and female mice was pooled. To drive Cre-dependent expression of channelrhodopsin2 (ChR2) in CA2 PCs for optogenetic experiments in acute hippocampal slices, we performed stereotactic injection (Nanoject II, Drummond #3-000-204) of Cre-dependent adenoassociated virus (rAAV5-EF1a-DIO-hChR2(E123T/T159C)-eYFP; UNC, lot AV4828b) targeted to CA1c, adjacent to the dorsal CA2 subfield in control and pilocarpine-treated Amigo2-Cre^+/-^ mice (male and female F1 generation progeny from a C57BL/6J x 129S1/SvlmJ cross). The stereotactic coordinates used were measured in millimeters relative to bregma (for anterior-posterior and medial-lateral coordinates, A-P and M-L) and relative to the surface of the brain (for dorsal-ventral coordinates, D-V) and were as follows: A-P -2.0, M-L ± 1.80, D-V -1.20.

For surgery, carprofen was given as an analgesic agent (5 mg/kg, subcutaneous, s.c.), and postoperative analgesia was carprofen-supplemented food (either 60 mL of MediGel CPF-74-05-5022 food gel, or Rodent MDs #MD150-2 2 mg carprofen tablets, each estimated to be a dose of approximately 5 mg/kg over 24 hours of consumption). Before the commencement of surgery animals were weighed to record baseline body weight and to calculate carprofen dose. Surgical procedures were performed under sterile conditions as approved by the Columbia University IACUC. Following surgery mice were group housed and allowed to recover for a minimum of two weeks to allow for viral expression before preparation of acute hippocampal slices or processing of brains for immunohistochemistry.

#### B. *Ex vivo* slice electrophysiology and data analysis

##### 1. Hippocampal slice preparation

Acute hippocampal slices were prepared using sucrose-substituted artificial cerebrospinal fluid (referred to as sucrose solution) and standard ACSF. ACSF and sucrose solutions were made with purified water that had been filtered through a 0.22 μm filter. ACSF contained (concentration expressed as millimolar, mM): 22.5 glucose, 125 NaCl, 25 NaHCO_3_, 2.5 KCl, 1.25 NaH_2_PO_4_, 3 sodium pyruvate, 1 ascorbic acid, 2 CaCl_2_, and 1 MgCl_2_. Sucrose solution used for slice preparation contained, in mM: 195 sucrose, 10 glucose, 25 NaHCO_3_, 2.5 KCl, 1.25 NaH_2_PO_4_, 2 sodium pyruvate, 0.5 CaCl_2_, 7 MgCl_2_. ACSF and sucrose cutting solution were prepared fresh before each experiment and the osmolarity was consistently measured to be between 315 and 325 mOsm. Sucrose solution was chilled on ice and bubbled with carbogen gas (95% O_2_ / 5% CO_2_) for at least 30 mins before slice preparation. A recovery beaker was prepared with a 50:50 mixture of ACSF and sucrose solution and warmed to approximately 32° C. Mice were deeply anesthetized by isoflurane inhalation and immediate incisions were made to sever the diaphragm and access the heart. Transcardial perfusion was performed with ice-cold carbogenated sucrose cutting solution for approximately 30 – 45 s. The mouse was decapitated and the brain quickly removed, at which point the hippocampi were dissected free, placed into an agar block, and secured to a vibratome slicing platform with cyanoacrylate adhesive. Hippocampal slices were cut at a thickness of 400 μm, parallel to the transverse plane. Slices were collected from the dorsal and intermediate hippocampus, as CA2 circuits have primarily been characterized within the dorsal hippocampus (Chevaleyre & Siegelbaum, 2010; Hitti & Siegelbaum, 2014; Kohara et al., 2014). Slices were transferred to the warmed recovery beaker and allowed to recover for 30 m, after which the beaker was allowed to come to room temperature and left for an additional 90 m.

##### 2. Whole-cell slice recordings

Recording and stimulation pipettes were prepared from borosilicate glass capillaries using a heated-filament puller programed to produce pipettes with a resistance of approximately 3 – 5 mOhm. Stimulation pipettes were filled with 1 M NaCl. Recording pipettes were filled with intracellular solution composed of, in mM: 135 potassium gluconate (C_6_H_11_KO_7_), 5 KCl, 0.2 EGTA (C_14_H_24_N_2_O_10_), 10 HEPES (C_8_H_18_N_2_O_4_S), 2 NaCl, 5 MgATP (C_10_H_16_N_5_O_13_P_3_ · Mg^2+^), 0.4 Na_2_GTP (C_10_H_16_N_5_O_14_P_3_ · Na^+^), 10 Na_2_ phosphocreatine (C_4_H_8_N_3_O_5_PNa_2_ · H_2_O), and biocytin (0.2% by weight). Intracellular solution was prepared on ice, the osmolarity was adjusted to approximately 295 mOsm, and the pH titrated to approximately 7.2. For voltage clamp recordings, intracellular solution instead contained cesium (Cs+) methanesulfonate (CH_3_O_3_SCs). Hippocampal slices were individually transferred to the recording chamber of an electrophysiology station where ACSF, warmed to 32° C and continually bubbled with carbogen, was perfused through the chamber at a rate of approximately 1 – 3 mL/min. Whole-cell recordings from CA2 and CA3 PCs was accomplished primarily through a “blind patching” approach, whereas CA1c_deep_ PCs were targeted by visual guidance to ensure cell position in the deep PCL. CA2 PCs were targeted using the end of the stratum lucidum (SL) as an anatomical landmark and cell identity was confirmed post-hoc based on cellular morphology as revealed by biocytin and colocalization with known CA2 markers, such as PCP4 (figure 1A, figure 2A). Cells were left unperturbed for several minutes before any recordings to ensure the membrane potential and seal were stable, and cells with a resting membrane potential more depolarized than -50 mV or a series resistance greater than 25 MΩ were discarded.

##### 3. Intrinsic properties

The resting membrane potential was measured by recording at least one minute in the current clamp configuration, without any injected current, and averaging the membrane potential. Current was then applied as needed to hold cells at -70 mV. Neuronal identity was assessed according to several physiological characteristics in the current clamp configuration: action potential firing patterns upon current injection, a low input resistance, and the absence of large spontaneous postsynaptic potentials clearly distinguished CA2 PCs from nearby CA3a PCs. Cell identity was confirmed by post-hoc microscopy (see below). Action potential firing patterns and a current-firing rate curve were measured using ten current steps, from +100 pA to +1000 pA (in 100 pA increments). Subthreshold membrane properties, namely the input resistance and membrane capacitance, were assessed using eleven current steps from -100 pA to +100 pA (in 20 pA increments). Voltage sag was measured using a series of negative current steps, and sag measured from whichever current step produced an initial hyperpolarization to between -100 mV and -105 mV. Voltage sag was expressed as a ratio between the peak hyperpolarization at the beginning of the pulse (ΔVm_peak_) and the difference between ΔVm_peak_ and the steady-state hyperpolarization (ΔVm_ss_) at the end of the current step: sag ratio = (ΔVm_peak_ - ΔVm_ss_) / ΔVm_peak_.

Rheobase current, action potential voltage threshold, afterhyperpolarization (AHP), and other action potential characteristics were assessed with a current ramp that reached 1000 pA over one second. The first action potential to result from the current ramp was isolated and a phase-plane plot constructed from the action potential waveform and the derivative of the membrane potential during the spike. Action potential onset was defined as the timepoint where the rate of depolarization exceeded 5 mV/ms. This timepoint was used to measure the rheobase current and the action potential voltage threshold. The phase-plane plot was used to measure to maximum rate of rise and rate of descent during the action potential. A gridded data interpolation was constructed from the raw action potential waveform, and this interpolant was used to measure the action potential half-width and the AHP. The AHP was defined as the difference between the action potential voltage threshold and the negative peak membrane potential occurring just after the action potential.

The voltage-dependent ramp of slow depolarization in CA2 PCs (slow ΔVm) was measured from membrane potential responses produced by the ten +100 pA current steps (the same membrane potential responses described above, used to measure the current-firing rate curve). For every cell, membrane potential was measured near the onset of the current step (initial Vm) and at the end of the current step (final Vm). The difference of the final and initial Vm was measured as the slow depolarization (slow ΔVm) that occurred over the one second duration of the current step. The slow ΔVm values from each cell were binned based on the corresponding initial Vm values into ten bins ranging from -68 mV to -48 mV, in two mV increments. In other words, the first bin collected slow ΔVm values from responses with an initial depolarization between -68 mV and -66 mV, the second bin from -66 mV to -64 mV, and so on. Membrane potential responses with an initial Vm more negative than -68 mV (or more positive than -48 mV) were discarded. In this way the amplitude of the slow depolarization could be directly compared across groups while controlling for the voltage dependence of the response. Analysis of intrinsic properties was performed in MATLAB and AxoGraph.

##### 4. Synaptic responses, optogenetics, and pharmacology

Electrical stimulation was delivered as a single 0.2 ms pulse (Digitimer Ltd. Constant voltage isolated stimulator, model DS2A-Mk.II) and photostimulation was delivered as a 2 ms pulse of 470 nm light produced by an LED light source (ThorLabs High-power 1-channel LED driver with pulse modulation, model DC2100). For input-output curves both electrical stimulation and photostimulation were delivered in ten steps of intensity, ranging from 8 V to 80 V (in 8 V increments) or from 10% to 100% light intensity (in 10% increments). Photostimulation trains consisted of fifteen 2 ms pulses delivered at 30 Hz, followed after 500 ms by an additional single 2 ms pulse. Postsynaptic potentials were measured in current clamp from an initial holding potential of -70 mV (except where specified as -55 mV). In the voltage clamp configuration cells were held at -75 mV (for EPSCs) or +10 mV (for IPSCs). The exponential time constant of decay tau was measured at the time point at which postsynaptic currents decayed from their peak amplitude to 36.8% of their peak (an approximation of 1/e). For each cell, only postsynaptic currents within a set range of amplitudes were pooled and their tau values averaged to yield one time constant per cell, such that comparisons of time constants between the control and SE groups utilized postsynaptic currents of comparable amplitudes. In recordings examining the DG input to CA2 PCs, optogenetic activation of granule cell mossy fiber axons occasionally resulted in polysynaptic responses (likely involving recruitment of CA3 PCs). Therefore cells were excluded from analysis if light-evoked postsynaptic responses occurred at a polysynaptic latency or involved multiple compounding potentials.

#### C. Anatomical Assessments

For immunohistochemistry mice were placed under deep isoflurane-induced anesthesia and transcardially perfused, first with 0.9% NaCl and then with 4% PFA. Brains were extracted and immersed whole in 4% PFA overnight at 4° C. The next day, brains were washed in 0.3% glycine in phosphate buffered saline (PBS) for one hour at room temperature on a shaker and were then rinsed three times briefly with PBS. Coronal or horizontal sections were prepared at a thickness of 60 μm using a vibratome. Post-hoc immunohistochemistry was performed by immersing 400-μm thick acute hippocampal slices in 4% paraformaldehyde (PFA) at the conclusion of recordings and fixing them overnight on a shaker at 4° C. Both 60 μm brain sections and 400 μm hippocampal slices were incubated for four hours at room temperature in PBS with 0.5% Triton-X. This solution was then exchanged for PBS containing primary antibodies and 0.1% Triton-X. The following antibodies were used: chicken anti-GFP (Aves #GFP-1020), rabbit anti-PCP4 (Sigma-Aldrich #HPA005792), mouse anti-RGS14 IgG2a (Neuromab #75-170), mouse anti-STEP IgG1 (Cell Signaling Technology #4396), and rabbit anti-CALB1 (Abcam #ab11426). Neuronal somata were visualized with a stain against the Nissl substance using the NeuroTrace 435/455 fluorescent dye (Invitrogen #N21479). Biocytin was visualized using streptavidin conjugated to Alexa Fluor 647 (Invitrogen #S21374). All images were acquired with a confocal microscope and processed using FIJI software.

### V. Statistical comparisons

All statistical tests were performed in GraphPad Prism or JASP. In all cases the cutoff for significance was set to a P value of 0.05. For parametric data, t-tests and ANOVA followed by Tukey-Kramer post hoc tests were used. For nonparametric data, Mann-Whitney U tests and Kruskal-Wallis followed by Dunn’s multiple comparisons test were used instead. Normality tests were conducted in Prism to confirm parametric statistics could be performed. Where variances were unequal, all data were log-transformed prior to analysis. In some cases of two-way ANOVA, the Geisser-Greenhouse correction and Holm’s or Sidak’s multiple comparisons test were used. Note that in cases where datasets had missing values, Prism applied a comparable mixed-effects model, which utilizes a compound symmetry covariance matrix fit using Restricted Maximum Likelihood (REML) with the Holm’s multiple comparisons test. Sample sizes were determined based on power analysis of pilot data using Instat2 (GraphPad).

## Acknowledgements

Supported by National Institutes of Health R01 NS106983, the American Epilepsy Society, and the New York State Office of Mental Health

---

**Supplementary Table 1.**
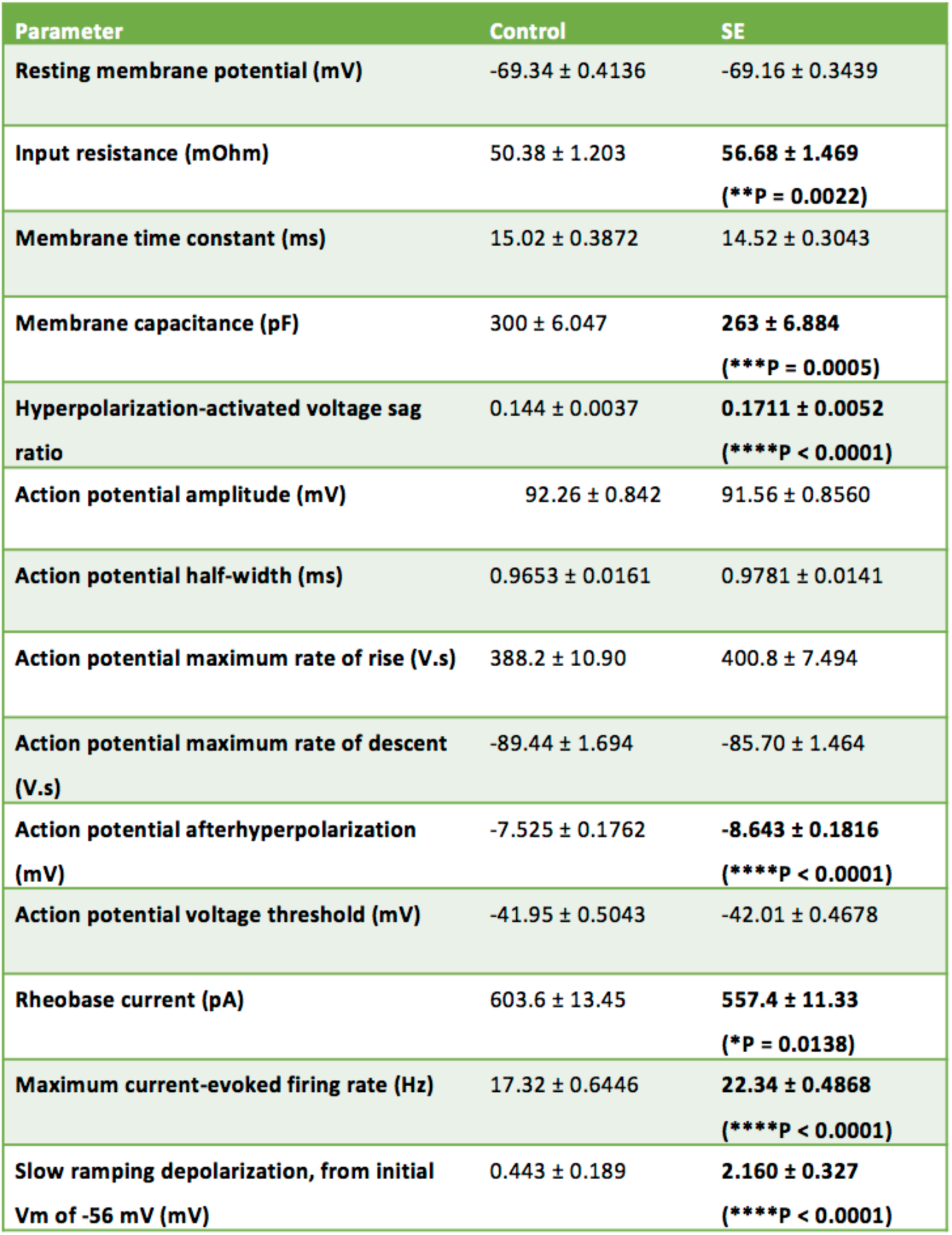
A summary of the intrinsic electrophysiological properties of CA2 PCs. Comparisons between CA2 PCs in hippocampal slices from control and SE mice revealed significant differences in several parameters. The membrane capacitance and rheobase current were decreased in Cells from SE mice, whereas the input resistance, maximum firing rate, hyperpolarization-activated voltage sag ratio, action potential afterhyperpolarization, and slow ramping depolarization were all increased in CA2 PCs from SE mice relative to controls.

| Parameter | Control | SE |
| --- | --- | --- |
| Resting membrane potential (mV) | -69.34 ± 0.4136 | -69.16 ± 0.3439 |
| Input resistance (mOhm) | 50.38 ± 1.203 | **56.68 ± 1.469**<br>(**P = 0.0022) |
| Membrane time constant (ms) | 15.02 ± 0.3872 | 14.52 ± 0.3043 |
| Membrane capacitance (pF) | 300 ± 6.047 | **263 ± 6.884**<br>(***P = 0.0005) |
| Hyperpolarization-activated voltage sag ratio | 0.144 ± 0.0037 | **0.1711 ± 0.0052**<br>(****P < 0.0001) |
| Action potential amplitude (mV) | 92.26 ± 0.842 | 91.56 ± 0.8560 |
| Action potential half-width (ms) | 0.9653 ± 0.0161 | 0.9781 ± 0.0141 |
| Action potential maximum rate of rise (V.s) | 388.2 ± 10.90 | 400.8 ± 7.494 |
| Action potential maximum rate of descent (V.s) | -89.44 ± 1.694 | -85.70 ± 1.464 |
| Action potential afterhyperpolarization (mV) | -7.525 ± 0.1762 | **-8.643 ± 0.1816**<br>(****P < 0.0001) |
| Action potential voltage threshold (mV) | -41.95 ± 0.5043 | -42.01 ± 0.4678 |
| Rheobase current (pA) | 603.6 ± 13.45 | **557.4 ± 11.33**<br>(*P = 0.0138) |
| Maximum current-evoked firing rate (Hz) | 17.32 ± 0.6446 | **22.34 ± 0.4868**<br>(****P < 0.0001) |
| Slow ramping depolarization, from initial Vm of -56 mV (mV) | 0.443 ± 0.189 | **2.160 ± 0.327**<br>(****P < 0.0001) |

**Supplementary Figure 1.**
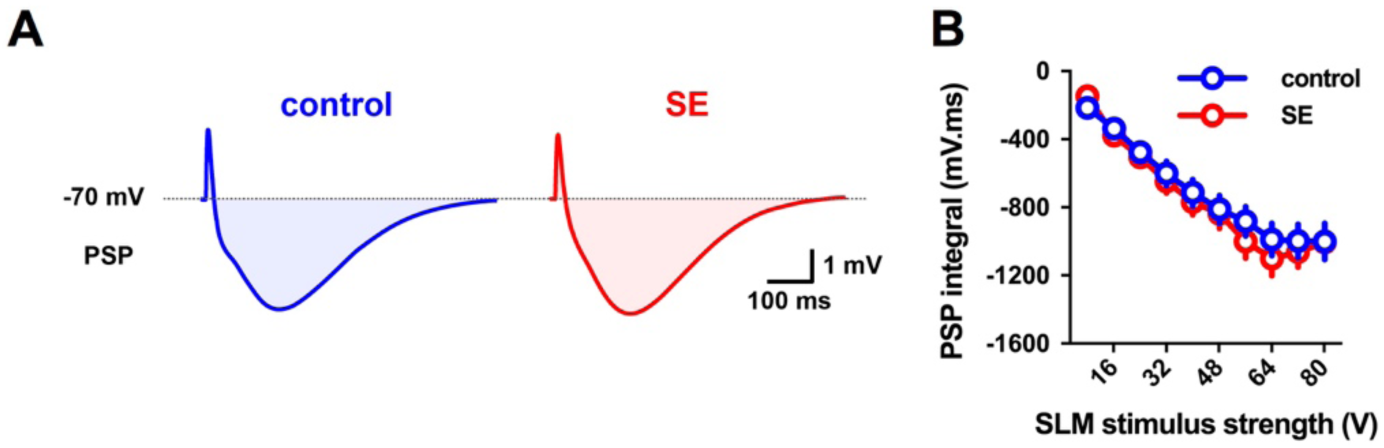
Inhibition in CA2 PCs elicited by activation of the EC LII inputs was not significantly altered in SE mice. **(A)** Representative averaged PSPs evoked by electrical stimulation in the SLM recorded in CA2 PCs from control (left, blue) and SE (right, red) mice. **(B)** The integral of the SLM-evoked PSP was not significantly different between the control and SE groups (mixed-effects model with Holm-Sidak’s test; P = 0.7934; n = 34 control, 31 SE).

**Supplementary Figure 2.**
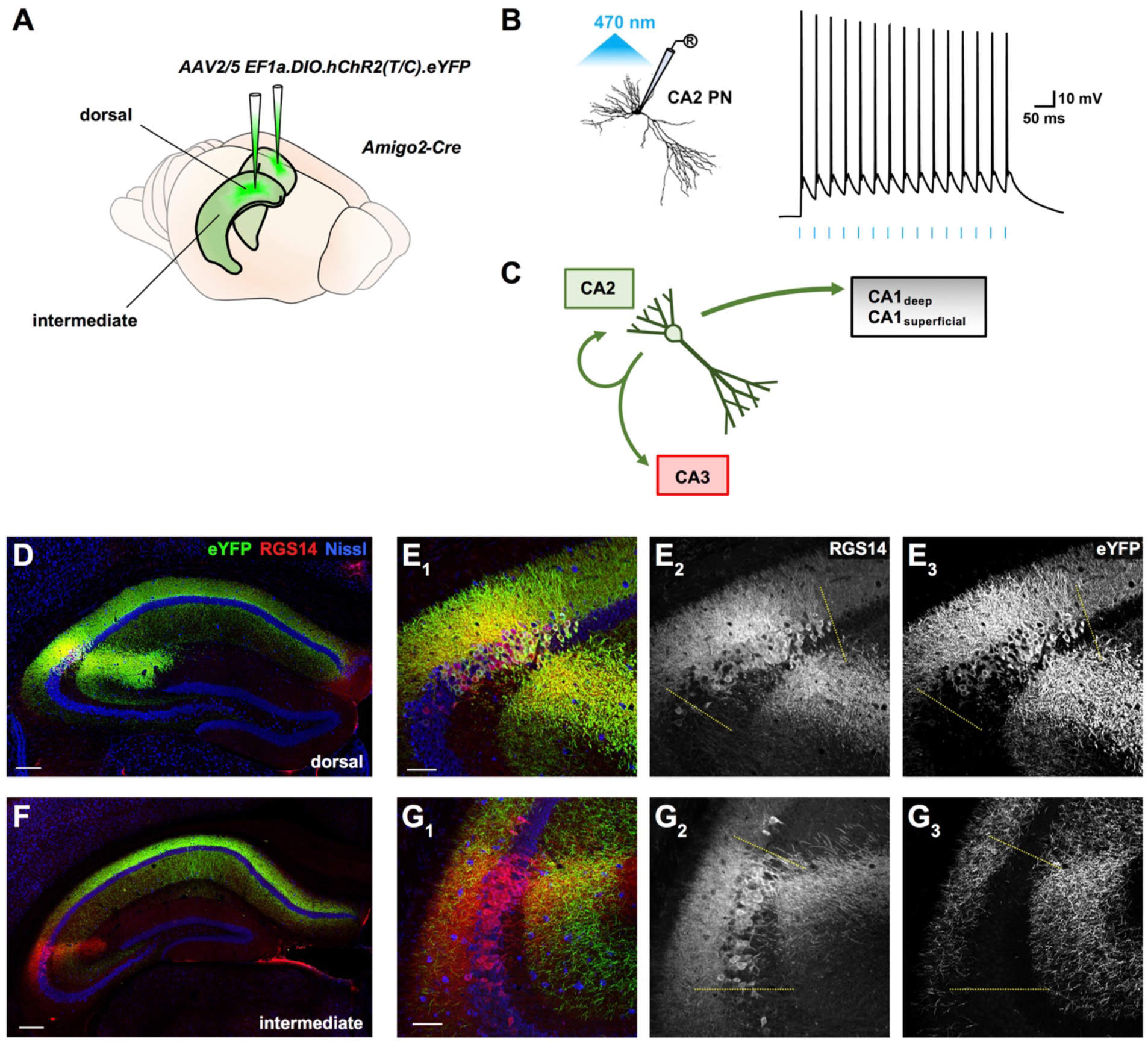
An optogenetic strategy using the Amigo2-Cre mouse line to investigate CA2 PC synaptic output in hippocampal slices from control and SE mice. **(A)** AAV was injected bilaterally into the dorsal hippocampus of Amigo2-Cre mice to drive channelrhodopsin-2 (ChR2-eYFP) expression in dorsal CA2 PCs (see Methods). **(B)** Representative current clamp recording from a CA2 PC illustrating the effect of photostimulation (blue lines) delivered in 2 ms pulses, 15 times at a frequency of 30 Hz. **(C)** A simplified circuit diagram of CA3–CA2–CA1 hippocampal circuit, in which CA2 PCs, like those in CA3, form both local recurrent excitatory axonal collaterals and projections to the CA1 region. CA2 preferentially excites CA1_deep_ PCs, and also back-projects to CA3. **(D)** A representative section from the dorsal hippocampus of AAV-injected Amigo2-Cre mice, showing ChR2-eYFP-expressing CA2 PCs near the site of viral injection. Scale bar is 200 μm. **(E_1_ - E_3_)** Immunohistochemistry showing the CA2 marker RGS14 (red) colocalized with ChR2-eYFP expression (green), demonstrating expression specificity in CA2 PCs. Yellow dashed lines indicate the approximate bounds of the CA2 subfield. Scale bar is 80 μm. **(F)** A representative section from the intermediate portion (approx. middle third) of the hippocampus from an AAV-ChR2-eYFP-injected Amigo2-Cre mouse, which contains longitudinally descending ChR2-eYFP-expressing CA2 axons originating from the dorsal hippocampus. Scale bar is 200 μm. **(G_1_ - G_3_)** In the intermediate hippocampus CA2 PCs (defined by RGS14 staining in red) did not express ChR2-eYFP expression but colocalized with ChR2-eYFP+ axon projections (green) from dorsal CA2 PCs. Yellow dashed lines indicate the approximate bounds of the CA2 subfield, identified by immunoreactivity for established selective markers of CA2 PCs (STEP, PCP4, or RGS14, see methods). Scale bar is 80 μm.

**Supplementary Figure 3.**
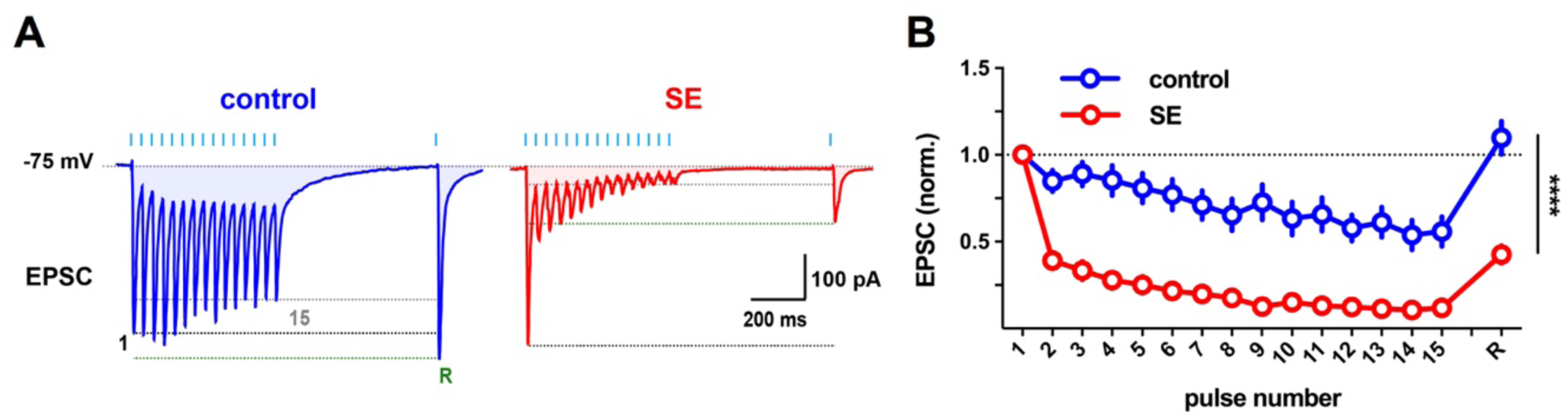
The granule cell mossy fiber input to CA2 PCs exhibited pronounced short-term depression following PILO-SE. **(A)** To examine short-term plasticity 15 light pulses (2 ms) were delivered at a frequency of 30 Hz, followed up by a single recovery pulse 500 ms after the train. Top, the blue lines indicate the timing of each 2 ms light pulse. Below, EPSCs in CA2 PCs from control and SE mice in response to the photostimulation train protocol. **(B)** In CA2 PCs from SE mice, the light-evoked EPSC exhibited a prominent short-term depression, which persisted to the recovery pulse.

**Supplementary Figure 4.**
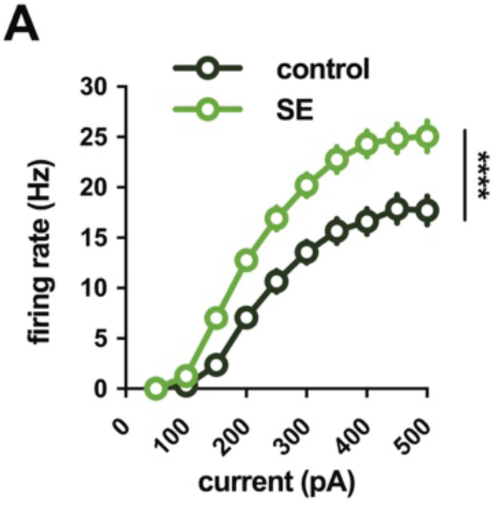
The intrinsic firing of CA1c_deep_ pyramidal cells is increased in slices from SE mice. **(A)** In slices from SE mice, CA1c_deep_ PNs fired action potentials at a higher rate upon direct current injection as compared to controls.

**Supplementary Figure 5.**
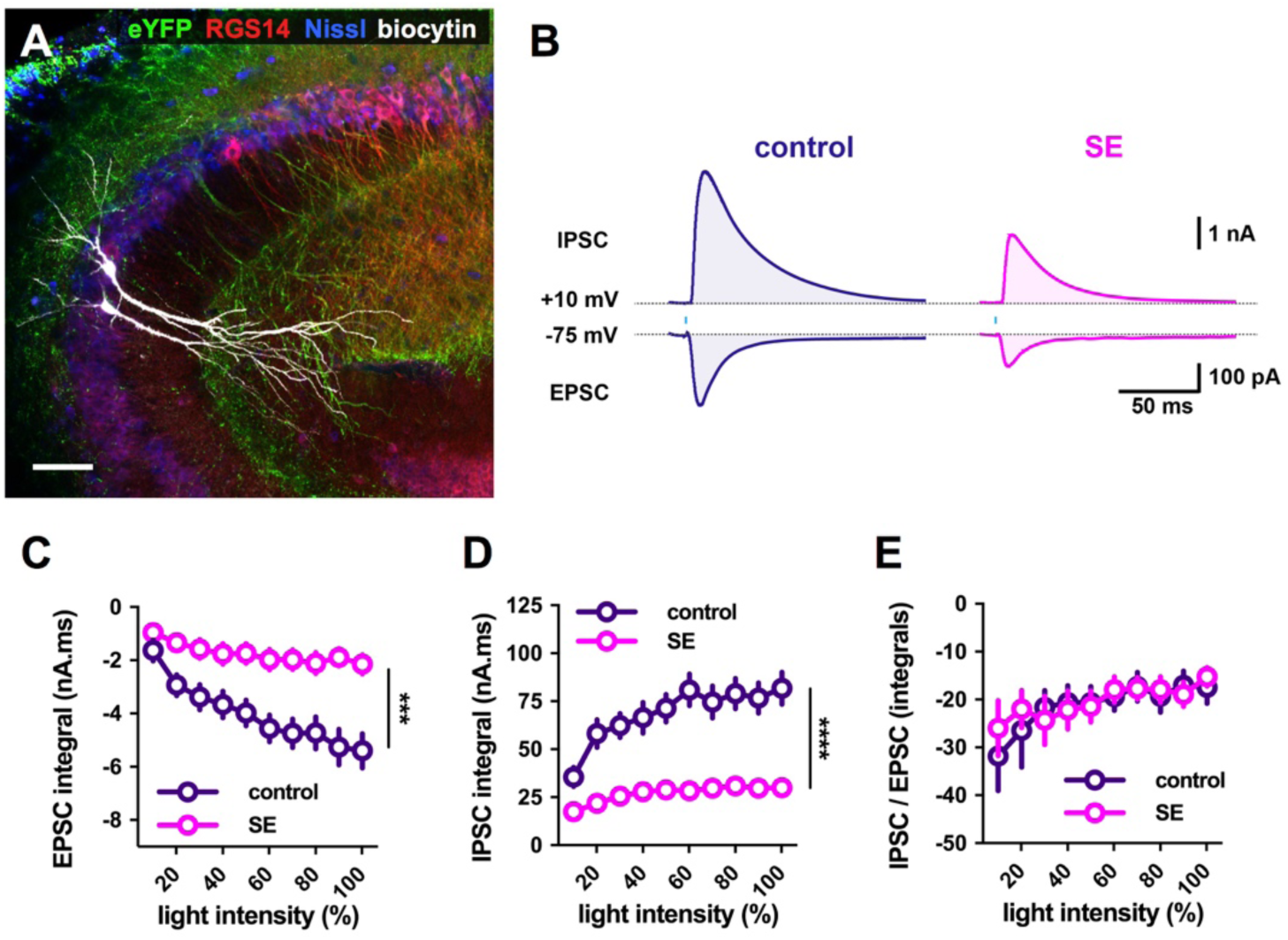
The CA2 back-projecting circuit to CA3a is weakened in slices from SE mice. **(A)** A representative image of two biocytin-filled CA3a PCs (white) surrounded by ChR2-eYFP-expressing CA2 axonal collaterals visible in the SO and SR (green), with CA2 PCs labeled using RGS14 (red) and neuronal somata labeled using a Nissl stain (blue). Scale bar is 80 μm. **(B)** Representative light-evoked averaged EPSCs and IPSCs recorded from CA3a PCs voltage clamped at -75 mV and +10 mV, respectively, in hippocampal slices from control (left, dark purple) and SE (right, magenta) mice. **(C)** The integral of the light-evoked EPSC was significantly reduced in CA3a PNs from SE mice (two-way ANOVA; ***P = 0.0004; n = 13 control, 19 SE). **(D)** The integral of the light-evoked IPSC was significantly reduced in CA3a PCs from SE mice (two-way ANOVA; ****P < 0.0001; n = 14 control, 19 SE). **(E)** The ratio of the integrals of the light-evoked IPSCs and EPSCs in CA3a PCs was not significantly altered in slices from SE mice (two-way ANOVA; P = 0.9059; n = 13 control, 19 SE).

**Supplementary Figure 6.**
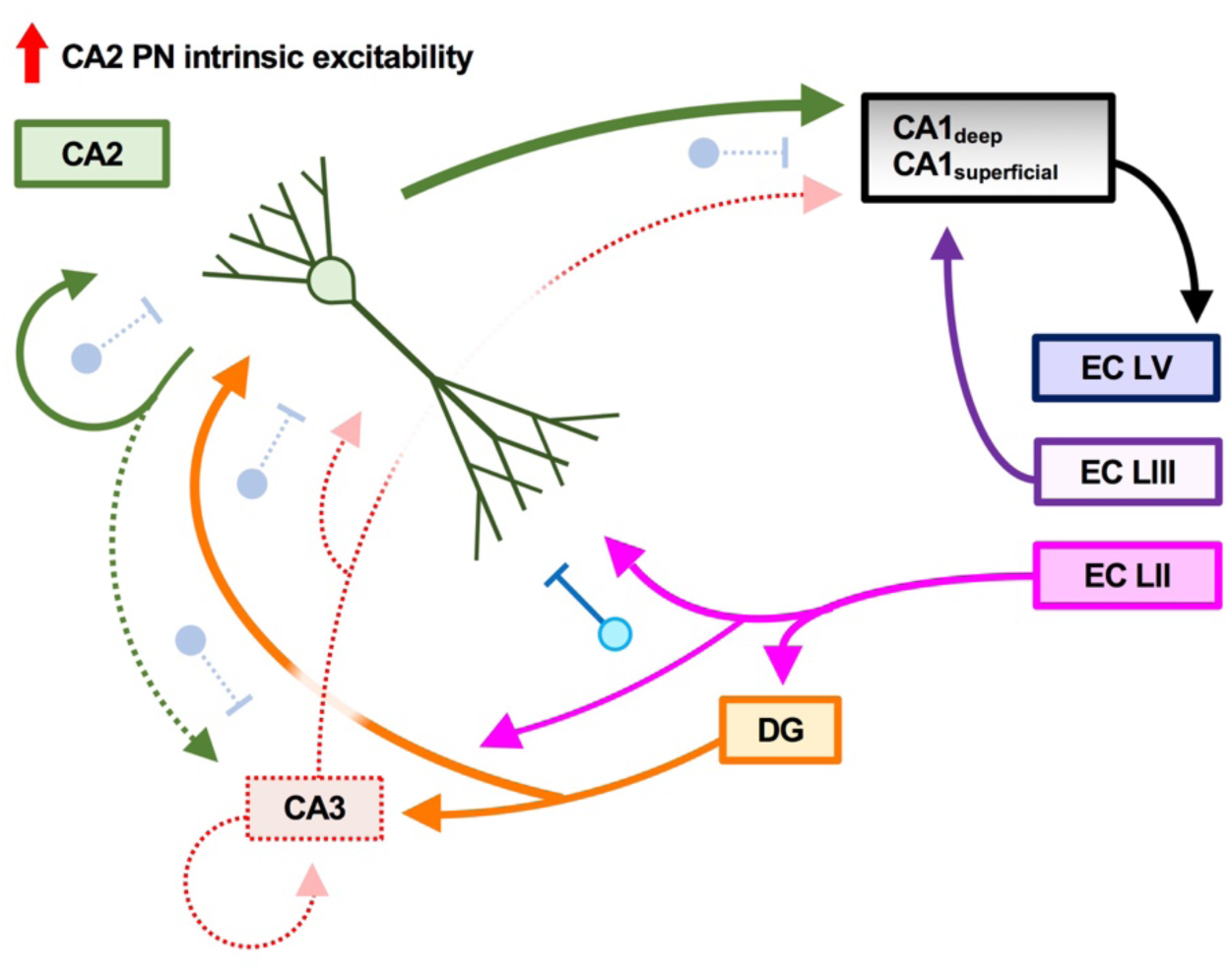
A summary of alterations to CA2 circuits following pilocarpine-induced status epilepticus. Reductions in GABAergic inhibition were observed in pathways examined, except for the entorhinal cortex projections in SLM to CA2. We found an increased strength of the excitatory mossy fiber inputs from DG granule cells to CA2 PCs and from CA2 PCs to CA1 PCs, as well as increased intrinsic excitability of CA2 and CA1 PCs.

**Supplemental Figure 7.**
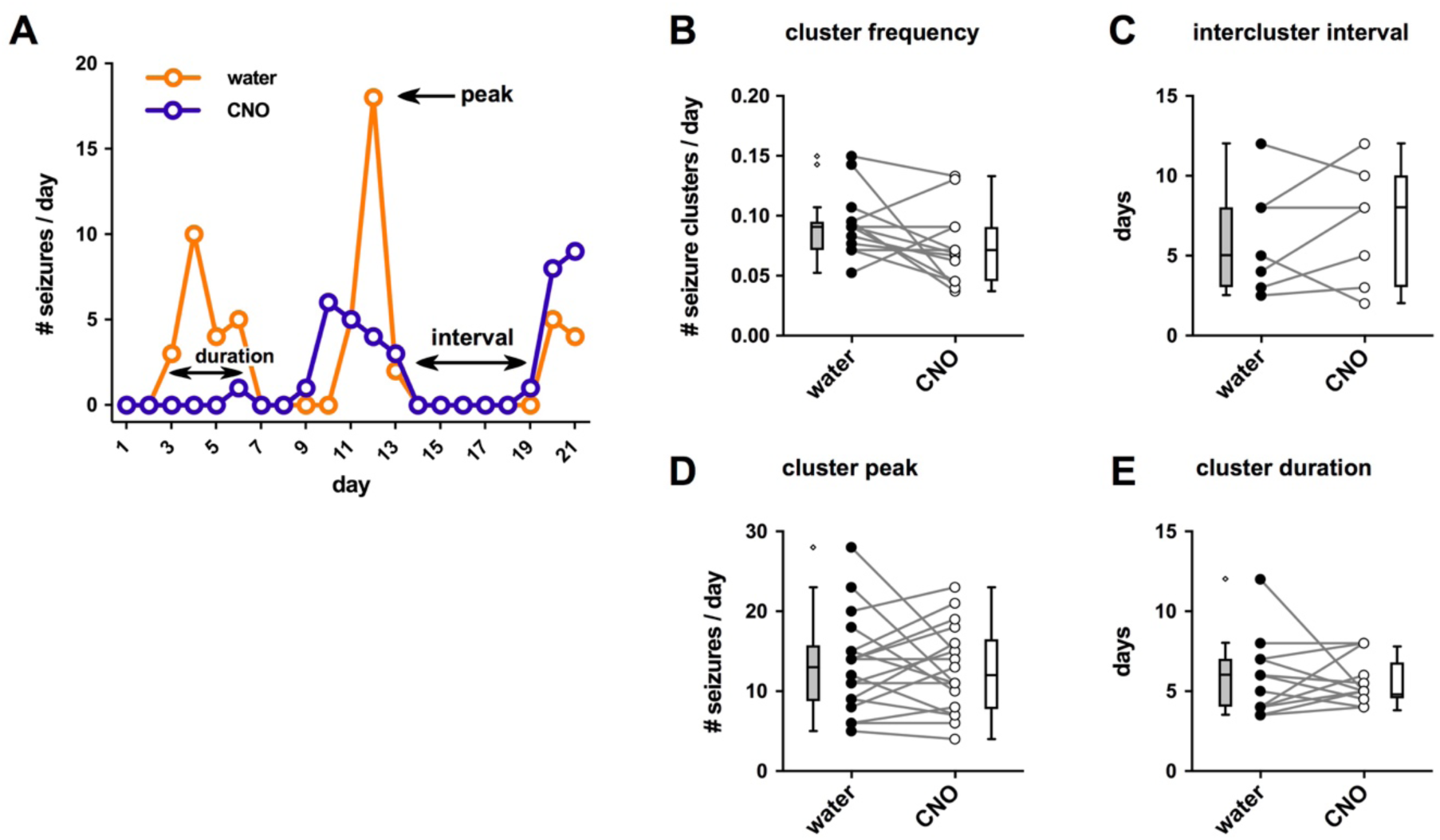
CA2 silencing did not alter seizure clustering. **(A)** A representative example of seizure clustering in one mouse expressing hM4D(Gi)-mCherry in CA2, with three weeks of daily seizure counts in absence of CNO (orange) and during delivery of CNO (purple). **(B)** The frequency of seizure clusters was not reduced by CNO treatment (paired t-test; t = 1.941, df = 14, *P* = 0.0727; n = 15). **(C)** The intercluster interval was not altered by CNO treatment (paired t-test; t = 0.7597, df = 6, *P* = 0.4762; n = 7). **(D)** The peak daily seizure total was not altered by CNO treatment (paired t-test; t = 0.3575, df = 17, *P* = 0.7251; n = 18). **(E)** The cluster duration was not altered by CNO treatment (paired t-test; t = 0.26731.941, df = 12, *P* = 0.7938; n = 13).

## REFERENCES

Alexander, G. M., Brown, L. Y., Farris, S., Lustberg, D., Pantazis, C., Gloss, B., … Dudek, S. M. (2018). CA2 neuronal activity controls hippocampal low gamma and ripple oscillations. ELife, 7, 1–25. https://doi.org/10.7554/eLife.38052

Andrioli, A., & Arellano, J. I. (2007). Quantitative analysis of parvalbumin-immunoreactive cells in the human epileptic hippocampus. Neuroscience, 149, 131–143. https://doi.org/10.1016/j.neuroscience.2007.07.029

Arnold, E. C., Mcmurray, C., Gray, R., & Johnston, D. (2019). Epilepsy-Induced Reduction in HCN Channel Expression Contributes to an Increased Excitability in Dorsal, But Not Ventral, Hippocampal CA1 Neurons. ENeuro, 6(April), 1–22.

Barmashenko, G., Hefft, S., Aertsen, A., Kirschstein, T., & Köhling, R. (2011). Positive shifts of the GABAA receptor reversal potential due to altered chloride homeostasis is widespread after status epilepticus. Epilepsia, 52(9), 1570–1578. https://doi.org/10.1111/j.1528-1167.2011.03247.x

Bartesaghi, R., Migliore, M., & Gessi, T. (2006). Input-output relations in the entorhinal cortex-dentate-hippocampal system: Evidence for a non-linear transfer of signals. Neuroscience, 142(1), 247–265. https://doi.org/10.1016/j.neuroscience.2006.06.001

Baud, M. O., Kleen, J. K., Mirro, E. A., Andrechak, J. C., King-Stephens, D., Chang, E. F., & Rao, V. R. (2018). Multi-day rhythms modulate seizure risk in epilepsy. Nature Communications, 9(1), 1–10. https://doi.org/10.1038/s41467-017-02577-y

Blümcke, I., Thom, M., Aronica, E., Armstrong, D. D., Bartolomei, F., Bernasconi, A., … Spreafico, R. (2013). International consensus classification of hippocampal sclerosis in temporal lobe epilepsy: A Task Force report from the ILAE Commission on Diagnostic Methods. Epilepsia, 54(7), 1315–1329. https://doi.org/10.1111/epi.12220

Boehringer, R., Polygalov, D., Huang, A. J. Y., Piskorowski, R. A., Chevaleyre, V., Mchugh, T. J., … Mchugh, T. J. (2017). Chronic Loss of CA2 Transmission Leads to Hippocampal Hyperexcitability. Neuron, 94(3), 642–655.e9. https://doi.org/10.1016/j.neuron.2017.04.014

Botterill, J. J., Lu, Y. L., LaFrancois, J. J., Bernstein, H. L., Alcantara-Gonzalez, D., Jain, S., … Scharfman, H. E. (2019). An Excitatory and Epileptogenic Effect of Dentate Gyrus Mossy Cells in a Mouse Model of Epilepsy. Cell Reports, 29(9), 2875–2889.e6. https://doi.org/10.1016/j.celrep.2019.10.100

Botterill, J. J., Vinod, K. Y., Gerencer, K. J., Teixeira, C. M., LaFrancois, J. J., & Scharfman, H. E. (2021). Bidirectional regulation of cognitive and anxiety-like behaviors by dentate gyrus mossy cells in male and female mice. Journal of Neuroscience, 41(11), 2475–2495. https://doi.org/10.1523/JNEUROSCI.1724-20.2021

Chevaleyre, V., & Siegelbaum, S. a. (2010). Strong CA2 pyramidal neuron synapses define a powerful disynaptic cortico-hippocampal loop. Neuron, 66(4), 560–572. https://doi.org/10.1016/j.neuron.2010.04.013

Cui, Z., Gerfen, C. R., & Young, W. S. (2013). Hypothalamic and other connections with dorsal CA2 area of the mouse hippocampus. Journal of Comparative Neurology, 521(8), 1844–1866. https://doi.org/10.1002/cne.23263

Du, F., Eid, T., Lothman, E. W., Kohler, C., & Schwarcz, R. (1995). Preferential neuronal loss in layer III of the medial entorhinal cortex in rat models of temporal lobe epilepsy. Journal of Neuroscience, 15(10), 6301–6313. https://doi.org/10.1523/jneurosci.15-10-06301.1995

Du, Fu, Whetsell, W. O., Abou-Khalil, B., Blumenkopf, B., Lothman, E. W., & Schwarcz, R. (1993). Preferential neuronal loss in layer III of the entorhinal cortex in patients with temporal lobe epilepsy. Epilepsy Research, 16(3), 223–233. https://doi.org/10.1016/0920-1211(93)90083-J

Freiman, T. M., Häussler, U., Zentner, J., Doostkam, S., Beck, J., Scheiwe, C., … Puhahn-schmeiser, B. (2021). Mossy fiber sprouting into the hippocampal region CA2 in patients with temporal lobe epilepsy. Hippocampus, 31(6), 1–13. https://doi.org/10.1002/hipo.23323

Goodkin, H. P., & Kapur, J. (2009). The impact of diazepam’s discovery on the treatment and understanding of status epilepticus. Epilepsia, 50(9), 2011–2018. https://doi.org/10.1111/j.1528-1167.2009.02257.x

Häussler, U., Rinas, K., Kilias, A., Egert, U., & Haas, C. A. (2016). Mossy fiber sprouting and pyramidal cell dispersion in the hippocampal CA2 region in a mouse model of temporal lobe epilepsy. Hippocampus, 26(5), 577–588. https://doi.org/10.1002/hipo.22543

He, H., Boehringer, R., Huang, A. J. Y., Overton, E. T. N., Polygalov, D., Okanoya, K., & Mchugh, T. J. (2021). CA2 inhibition reduces the precision of hippocampal assembly reactivation. Neuron, 109, 1–14. https://doi.org/10.1016/j.neuron.2021.08.034

Hendricks, W. D., Chen, Y., Bensen, A. L., Westbrook, G. L., & Schnell, E. (2017). Short-term depression of sprouted mossy fiber synapses from adult-born granule cells. Journal of Neuroscience, 37(23), 5722–5735. https://doi.org/10.1523/JNEUROSCI.0761-17.2017

Hendricks, W. D., Westbrook, G. L., & Schnell, E. (2019). Early detonation by sprouted mossy fibers enables aberrant dentate network activity. Proceedings of the National Academy of Sciences of the United States of America, 166(22), 10994–10999. https://doi.org/10.1073/pnas.1821227116

Hitti, F. L., & Siegelbaum, S. a. (2014). The hippocampal CA2 region is essential for social memory. Nature, 508(7494), 88–92. https://doi.org/10.1038/nature13028

Iyengar, S. S., LaFrancois, J. J., Friedman, D., Drew, L. J., Denny, C. A., Burghardt, N. S., … Scharfman, H. E. (2015). Suppression of adult neurogenesis increases the acute effects of kainic acid. Experimental Neurology, 264, 135–149. https://doi.org/10.1016/j.expneurol.2014.11.009

Jaffe, D. B., & Brenner, R. (2018). A computational model for how the fast afterhyperpolarization paradoxically increases gain in regularly firing neurons. Journal of Neurophysiology, 119(4), 1506–1520. https://doi.org/10.1152/jn.00385.2017

Jain, S., LaFrancois, J. J., Botterill, J. J., Alcantara-Gonzalez, D., & Scharfman, H. E. (2019). Adult neurogenesis in the mouse dentate gyrus protects the hippocampus from neuronal injury following severe seizures. Hippocampus, 29(8), 1–27. https://doi.org/10.1002/hipo.23062

Janz, P., Savanthrapadian, S., Häussler, U., Kilias, A., Nestel, S., Kretz, O., … Haas, C. A. (2017). Synaptic Remodeling of Entorhinal Input Contributes to an Aberrant Hippocampal Network in Temporal Lobe Epilepsy. Cerebral Cortex, 27(3), 2348–2364. https://doi.org/10.1093/cercor/bhw093

Kay, K., & Frank, L. M. (2019). Three brain states in the hippocampus and cortex. Hippocampus, 29(3), 184–238. https://doi.org/10.1002/hipo.22956

Kohara, K., Pignatelli, M., Rivest, A. J., Jung, H.-Y., Kitamura, T., Suh, J., … Tonegawa, S. (2014). Cell type-specific genetic and optogenetic tools reveal hippocampal CA2 circuits. Nature Neuroscience, 17(2), 269–279. https://doi.org/10.1038/nn.3614

Krook-Magnuson, E., Armstrong, C., Bui, A., Lew, S., Oijala, M., & Soltesz, I. (2015). In vivo evaluation of the dentate gate theory in epilepsy. The Journal of Physiology, 10, 2379–2388. https://doi.org/10.1113/JP270056

Kwan, P., & Sander, J. W. (2004). The natural history of epilepsy: An epidemiological view. Journal of Neurology, Neurosurgery and Psychiatry, 75(10), 1376–1381. https://doi.org/10.1136/jnnp.2004.045690

Kwan, Patrick, & Brodie, M. J. (2000). Early Identification of Refractory Epilepsy. The New England Journal of Medicine, 342(5), 314–319.

Lehr, A. B., Kumar, A., Tetzlaff, C., Hafting, T., Fyhn, M., & St, T. M. (2021). CA2 beyond social memory : Evidence for a fundamental role in hippocampal information processing. Neuroscience and Biobehavioral Reviews, 126(March), 398–412. https://doi.org/10.1016/j.neubiorev.2021.03.020

MacLaren, D. A. A., Browne, R. W., Shaw, J. K., Radhakrishnan, S. K., Khare, P., España, R. A., & Clark, S. D. (2016). Clozapine N-oxide administration produces behavioral effects in long-evans rats: Implications for designing DREADD experiments. ENeuro, 3(5). https://doi.org/10.1523/ENEURO.0219-16.2016

Madisen, L., Mao, T., Koch, H., Zhuo, J. M., Berenyi, A., Fujisawa, S., … Zeng, H. (2012). A toolbox of Cre-dependent optogenetic transgenic mice for light-induced activation and silencing. Nature Neuroscience, 15(5), 793–802. https://doi.org/10.1038/nn.3078

Mahler, S. V., & Aston-Jones, G. (2018). CNO Evil? Considerations for the Use of DREADDs in Behavioral Neuroscience. Neuropsychopharmacology, 43(5), 934–936. https://doi.org/10.1038/npp.2017.299

McHugh, T. J., Jones, M. W., Quinn, J. J., Balthasar, N., Coppari, R., Elmquist, J. K., … Tonegawa, S. (2007). Dentate Gyrus NMDA Receptors Mediate Rapid Pattern Separation in the Hippocampal Network. Science, 317(July), 94–99. https://doi.org/10.4324/9781315657455-44

Meira, T., Leroy, F., Buss, E. W., Oliva, A., Park, J., & Siegelbaum, S. A. (2018). A hippocampal circuit linking dorsal CA2 to ventral CA1 critical for social memory dynamics. Nature Communications, 9(4163), 1–14. https://doi.org/10.1038/s41467-018-06501-w

Nasrallah, K., Therreau, L., Robert, V., Huang, A. J. Y., Mchugh, T. J., Piskorowski, R. A., … Mchugh, T. J. (2019). Routing Hippocampal Information Flow through Parvalbumin Interneuron Plasticity in Area CA2. Cell Reports, 27(1), 86–98.e3. https://doi.org/10.1016/j.celrep.2019.03.014

Ogren, J. A., Bragin, A., Wilson, C. L., Hoftman, G. D., Lin, J. J., Dutton, R. A., … Staba, R. J. (2009). Three-dimensional hippocampal atrophy maps distinguish two common temporal lobe seizure-onset patterns. Epilepsia, 50(6), 1361–1370. https://doi.org/10.1111/j.1528-1167.2008.01881.x

Okamoto, K., & Ikegaya, Y. (2018). Recurrent connections between CA2 pyramidal cells. Hippocampus, (May), 321513. https://doi.org/10.1101/321513

Okuyama, T., Kitamura, T., Roy, D. S., Itohara, S., & Tonegawa, S. (2016). Ventral CA1 neurons store social memory. Science, 353(6307), 1536–1541.

Oliva, A., Fernández-Ruiz, A., Buzsáki, G., & Berényi, A. (2016). Role of Hippocampal CA2 Region in Triggering Sharp-Wave Ripples. Neuron, 91, 1–14. https://doi.org/10.1016/j.neuron.2016.08.008

Oliva, A., Fernández-Ruiz, A., Leroy, F., & Siegelbaum, S. A. (2020). Hippocampal CA2 sharp-wave ripples reactivate and promote social memory. Nature, (November 2019). https://doi.org/10.1101/2020.03.03.975599

Racine, R. J. (1972). Modification of Seizure Activity by Electrical Stimulation: II. Motor Seizure. Electroencephalography and Clinical Neurophysiology, 32, 281–294.

Scharfman, H. E. (2019). The Dentate Gyrus and Temporal Lobe Epilepsy: An “Exciting” Era. Epilepsy Currents, 19(4), 1–7. https://doi.org/10.1177/1535759719855952

Schmeiser, B., Zentner, J., Prinz, M., Brandt, A., & Freiman, T. M. (2017). Extent of mossy fiber sprouting in patients with mesiotemporal lobe epilepsy correlates with neuronal cell loss and granule cell dispersion. Epilepsy Research, 129, 51–58. https://doi.org/10.1016/j.eplepsyres.2016.11.011

Smith, Z. Z., Benison, A. M., Bercum, F. M., Dudek, F. E., & Barth, D. S. (2018). Progression of convulsive and nonconvulsive seizures during epileptogenesis after pilocarpine-induced status epilepticus. Journal of Neurophysiology, 119, 1818–1835. https://doi.org/10.1152/jn.00721.2017

Spencer, S. S. (2002). Neural networks in human epilepsy: Evidence of and implications for treatment. Epilepsia, 43(3), 219–227. https://doi.org/10.1046/j.1528-1157.2002.26901.x

Srinivas, K. V, Buss, E. W., Sun, X. Q., Santoro, B., Takahashi, H., Nicholson, D. A., & Siegelbaum, S. A. (2017). The Dendrites of CA2 and CA1 Pyramidal Neurons Differentially Regulate Information Flow in the Cortico-Hippocampal Circuit. Journal of Neuroscience, 37(12), 3276–3293. https://doi.org/10.1523/JNEUROSCI.2219-16.2017

Stegen, M., Kirchheim, F., Hanuschkin, A., Staszewski, O., Veh, R. W., & Wolfart, J. (2012). Adaptive intrinsic plasticity in human dentate gyrus granule cells during temporal lobe epilepsy. Cerebral Cortex, 22(9), 2087–2101. https://doi.org/10.1093/cercor/bhr294

Steve, T. A., Jirsch, J. D., & Gross, D. W. (2014). Quantification of subfield pathology in hippocampal sclerosis: A systematic review and meta-analysis. Epilepsy Research, 108(8), 1279–1285. https://doi.org/10.1016/j.eplepsyres.2014.07.003

Sun, Q., Sotayo, A., Cazzulino, A. S., Snyder, A. M., Denny, C. A., Siegelbaum, S. A., … Denny, C. A. (2017). Proximodistal Heterogeneity of Hippocampal CA3 Pyramidal Neuron Intrinsic Properties, Connectivity, and Reactivation during Memory Recall. Neuron, 95(3), 656–672.e3. https://doi.org/10.1016/j.neuron.2017.07.012

Tamamaki, N., Abe, K., & Nojyo, Y. (1988). Three-dimensional analysis of the whole axonal arbors originating from single CA2 pyramidal neurons in the rat hippocampus with the aid of a computer graphic technique. Brain Research, 452(1–2), 255–272. https://doi.org/10.1016/0006-8993(88)90030-3

Tolner, E. A., Frahm, C., Metzger, R., Gorter, J. A., Witte, O. W., Lopes da Silva, F. H., & Heinemann, U. (2007). Synaptic responses in superficial layers of medial entorhinal cortex from rats with kainate-induced epilepsy. Neurobiology of Disease, 26(2), 419–438. https://doi.org/10.1016/j.nbd.2007.01.009

Valero, M., Cid, E., Averkin, R. G., Aguilar, J., Sanchez-Aguilera, A., Viney, T. J., … de la Prida, L. M. (2015). Determinants of different deep and superficial CA1 pyramidal cell dynamics during sharp-wave ripples. Nature Neuroscience, 18(9), 1281–1290. https://doi.org/10.1038/nn.4074

Williamson, A., & Spencer, D. D. (1994). Electrophysiological characterization of CA2 pyramidal cells from epileptic humans. Hippocampus, 4(2), 226–237.

Winawer, M. R., Makarenko, N., McCloskey, D. P., Hintz, T. M., Nair, N., Palmer, A. A., & Scharfman, H. E. (2007). Acute and chronic responses to the convulsant pilocarpine in DBA/2J and A/J mice. Neuroscience, 149(2), 465–475. https://doi.org/10.1016/j.neuroscience.2007.06.009

Wittner, L., Huberfeld, G., Clemenceau, S., Eross, L., Dezamis, E., Entz, L., … Miles, R. (2009). The epileptic human hippocampal cornu ammonis 2 region generates spontaneous interictal-like activity in vitro. Brain, 132(11), 3032–3046. https://doi.org/10.1093/brain/awp238

Wolfart, J., & Laker, D. (2015). Homeostasis or channelopathy ? Acquired cell type-specific ion channel changes in temporal lobe epilepsy and their antiepileptic potential. Frontiers in Physiology, 6(June), 1–23. https://doi.org/10.3389/fphys.2015.00168

Wozny, C., Gabriel, S., Jandova, K., Schulze, K., Heinemann, U., & Behr, J. (2005). Entorhinal cortex entrains epileptiform activity in CA1 in pilocarpine-treated rats. Neurobiology of Disease, 19(3), 451–460. https://doi.org/10.1016/j.nbd.2005.01.016

Wyeth, M., Nagendran, M., & Buckmaster, P. S. (2020). Ictal onset sites and γ-aminobutyric acidergic neuron loss in epileptic pilocarpine-treated rats. Epilepsia, 61(5), 856–867. https://doi.org/10.1111/epi.16490

